# Not All Butterflies Are Monarchs: Compass Systems in the Red Admiral (*Vanessa atalanta*), a European Diurnal Migrant

**DOI:** 10.1101/2025.05.01.651646

**Authors:** Aleksandr Pakhomov, Anatoly Shapoval, Nazar Shapoval, Dmitry Kishkinev

## Abstract

Seasonal migration in animals is a widespread and complex phenomenon, yet the mechanisms underlying orientation and navigation remain poorly understood in many taxa. While significant progress has been made in migratory birds, where multiple compass systems are well described, similar knowledge for migratory Lepidoptera remains limited. Most insights into butterfly and moth orientation come from just two species: the monarch butterfly (*Danaus plexippus*) and the Australian Bogong moth (*Agrotis infusa*), both of which possess multimodal compass systems involving visual and geomagnetic cues. However, these species follow unusual migratory strategies that include diapause or aestivation and multigenerational round-trip migrations between breeding region and very specific non-breeding areas, that are not representative of most other Lepidoptera migrants. In contrast, European species such as the painted lady (*Vanessa cardui*) and red admiral (*Vanessa atalanta*) undertake regular seasonal migrations involving multiple generations and no diapause, yet the sensory mechanisms guiding their orientation remain largely unexplored.

Here, we report findings from a two-year study investigating compass orientation in red admirals using a flight simulator under a range of controlled light and magnetic conditions. Our experiments yielded three key results: (1) red admirals orient using solar cues when selecting migratory direction; (2) the sun compass in this species appears to be time-independent, as clock-shifted individuals did not alter orientation; and (3) there is no sign of magnetic sense in red admirals. These findings sharply contrast with those from monarch butterflies, which rely on both time-compensated sun and light-dependent magnetic compasses. Our results reveal important interspecific variation in compass use among diurnal Lepidoptera and underscore the need to expand orientation research beyond traditional model systems to better understand the diversity of migratory strategies and orientation mechanisms in migratory Lepidoptera.

## Introduction

Each year, billions of animals undertake seasonal migrations, embarking on journeys that span vast distances and encompass a wide range of environments. These remarkable movements, observed across diverse taxa (from tiny insects to large mammals) represent one of nature’s most compelling phenomena. Understanding how animals accomplish these feats has long been a central focus in behavioural ecology, particularly the mechanisms underlying orientation (the ability to determine and maintain a direction of migration using compasses) and navigation (the capacity to determine one’s position relative to a goal without direct sensory contact using a map). To date, our understanding of these mechanisms is largely based on research involving a limited number of model taxa, most notably migratory birds. In birds, for example, at least three independent compass systems have been identified. The first is the light-dependent magnetic compass, which allows birds to detect the inclination angle of the Earth’s magnetic field (Wiltschko & Wiltschko, 1972). The second and third are astronomical compasses, based on celestial cues: the star compass (Emlen, 1967; Emlen, 1970) and the sun compass (Kramer, 1953; Able, 1982; Schmidt-Koenig, 1990).

In contrast to migratory birds, there remains a significant knowledge gap concerning the orientation mechanisms of migratory Lepidoptera. Following the invention of the flight simulator— a specialised setup for studying orientation in insects—in the early 2000s (Mouritsen & Frost, 2002), there has been a substantial increase in studies investigating the sensory basis of these processes in butterflies and moths. However, most of what we know about the compass systems of these small creatures come from studies on just two species: the North American monarch butterfly (*Danaus plexippus*, Linnaeus, 1758) and the Australian Bogong moth (*Agrotis infusa*). Monarch butterflies are the diurnal migrants and each autumn undertake 4000 km journey from the US and southern Canada to overwinter in the oyamel fir forests in central Mexico (Urquhart and Urquhart, 1978). They mostly rely on information from the sun using an antenna-based time-dependent sun compass system for orientation (Perez et al., 1997; Mouritsen and Frost, 2002; Reppert et al., 2004; Stalleicken et al., 2005; Merlin et al., 2009; Guerra et al., 2012) and coldness triggers the northward flight direction in spring (Guerra and Reppert, 2013). Studies suggested that this species possess the light-dependent inclination magnetic compass similar to avian magnetic compass, which requires UV-A/blue lights with specific wavelengths (Guerra et al., 2014; Wan et al., 2021; Kendzel et al., 2023) but it functions as a backup compass system: after clock-shift treatments, monarchs butterflies fly in the predicted, adjusted orientation despite being exposed to the Earth’s magnetic field and may only use the magnetic compass when daylight sky cues are unavailable (Reppert and de Roode, 2018). However, unlike the sun compass, there is no consensus among researchers regarding the existence of the magnetic compass in monarchs due to contardictory results obtained in other studies. Monarch butterflies did not show significant group-level season-appropriate orientation under simulated overcast conditions (Mouritsen and Frost, 2002; Stalleicken et al., 2005; Green et al., 2024). Nonetheless, this may be attributed to the absence of UV light with wavelengths between 380 and 420 nm, which was blocked by the diffuser materials used in these studies (Guerra et al., 2014; Green et al., 2024).

Another well-known Lepidopteran migrant, the Australian Bogong moth, undertakes a 1000 km journey each austral spring from various regions in southeast Australia to mountain caves in the alpine areas of New South Wales and Victoria (Warrant et al., 2016). It has been shown that these moths steer their nocturnal migratory flight by using information from the geomagnetic field and correlating it with visual landmarks (Dreyer et al., 2018a). Moreover, according to results of more recent study, the Bolong moth can use information from the stellar sky to choose appropriate migratory direction (Dreyer et al., 2024): under naturalistic moonless night skies and in a nulled geomagnetic field (no magnetic compass information), moths were oriented in their seasonally appropriate migratory directions. Additionally, they flew in a similar migratory direction under the same magnetic field condition when exposed to an artificial stellar sky simulating the sky of the study site, and accordingly changed their orientation when this artificial sky was rotated by 180°.

Both monarch butterflies and Bogong moths share common migratory features: they undertake regular annual multigenerational migrations, where the *same* individuals travel from breeding grounds to specific non-breeding sites and then begin the return migration in the following season. After spending several months in diapause—a hormonally controlled state of dormancy that helps monarch butterflies survive the winter—or in a hibernation-like ‘summer sleep’ called aestivation, which enables moths to endure hot summer conditions, they begin their return migration to their original locations to mate, lay eggs, and die. Subsequent generations either continue northward to recolonise mid-summer breeding areas (monarchs) or prepare for the next migratory season (moths). However, this migratory strategy is not widespread among many other migratory Lepidoptera (Reppert and de Roode, 2018), which typically exhibit year-round, continuous movements without entering a diapause phase. These migrations involve multiple generations per year, as seen in species such as the painted lady (*Vanessa cardui*) and red admiral (*Vanessa atalanta*) in Europe. Every year painted lady butterflies perform their incredible, multigenerational annual round-trip of about 12,000 to 14,000 km between tropical West Africa and Europe (Talavera and Vila, 2016; Talavera et al., 2018; Talavera et al., 2023; Reich et al., 2024) and even over 4000 km trans-oceanic flight between Africa nd South America (Suchan et al., 2024), strongly suggestiong that well-developed compass systems are at play. Red admirals undertake a fairly regular migration from northern Europe to the Mediterranean region in autumn (Williams, 1951; Benvenuti et al., 1994; Stefanescu, 2001; Brattström et al., 2018; Pakhomov et al., 2023). Unlike the North American monarch butterfly, red admirals do not hibernate upon arrival; instead, they overwinter in the Mediterranean region, where they lay eggs on nettles (*Urtica spp.)* (Stefanescu, 2001). In spring, the new generation emerges and begins migrating northwards, where they lay eggs again on host plants. One or two additional summer generations then continue recolonisation of northern Europe (Stefanescu, 2001) and in autumn, the main wave of red admirals starts their southward migration towards wintering sites in southern Europe and North Africa. Due to this full annual cycle, red admirals show two peaks of abundance in different parts of Europe, with their numbers in autumn being dramatically higher than in spring (Stefanescu, 2001; Cuadrado, 2017; Pakhomov et al., 2023).

Although these species are widely studied and considered as popular models in butterfly migration research, only a couple of studies have attempted to investigate their orientation abilities under lab-controlled conditions (Brattström, 2007; Nesbit et al., 2009), along with European migratory moths (Dreyer et al., 2018b). Our study endeavours to fill the knowledge gap of compass systems in European migratory butterflies: we conducted orientation experiments on red admirals tested in a flight simulator under different light and magnetic field conditions during spring and autumn migrations. We hypothesised that red admirals, like other diurnal Lepidopteran migrants such as monarch butterflies, primarily rely on visual cues (the sun compass) and compensate for the sun’s daily movement using internal clocks. Further, we investigated whether they are able to choose an appropriate migratory direction when access to the sun and sun-based cues is restricted, and the geomagnetic field is the only available directional cue.

## Materials and Methods

### Butterflies’ capture and husbandry

Red admirals of both sexes were collected during spring (May 2024) and autumn migration seasons (August-September, 2023-2024), using large “Rybachy-type” traps (Figure S1, see details in Brattström et al., 2018 and Pakhomov et al., 2023) at the “Fringilla” field station (Kaliningrad region, Russia). Upon capture, all butterflies were housed indoors in a windowless laboratory chamber, within large mesh cages (47 × 47 × 47 cm). They were kept under controlled conditions, including a constant temperature (21–22 °C during the light phase and 15-17 °C during the dark phase), 75–80% relative humidity, and an artificial photoperiod mimicking natural day-night cycles. SunLike LED bulbs (as a main light source) and ZooDa Bird Compact bulbs (as a UV source) were automatically switched on/off at local sunrise/sunset by the smart dimmer Shelly 4Pro (Allterco Robotics, Bulgaria). The butterflies were fed with 10% honey solution (in water) and were housed under such condition for at least 3-5 days before testing to recover from stress induced by capture. After orientation tests, all red admirals were released in good body condition well before the migration of their conspecifics was finished. All animal procedures were approved by the Ethics Committee of the Zoological Institute of the Russian Academy of Sciences (Permit No. 1-15/24-03-2023).

### Flight simulator

All behavioural experiments were performed in flight simulators modified from Mouritsen and Frost (2002) which allow to obtain butterfly flight tracks and analyse orientation. The flight simulator was made of non-magnetic materials (PVC, aluminium, PETG) and comprised three main components (Figure S1B,C):

1. **a flight chamber**: a white plastic cylinder (diameter 45 cm, height 50 cm) placed vertically on a plastic/aluminium table. A 120-mm diameter 3D-printed plastic pipe, with hundreds of 3-mm holes, was positioned at the center of the table to create laminar airflow (≤ 1 m/c). This airflow was generated by a PWM-controlled computer fan. The airflow was required to stimulate a tethered butterfly to engage in active flight (Mouritsen and Frost, 2002);
2. **a butterfly attachment apparatus** consisting of a plastic bar on the top of the cylinder with an encoder at its center. The 15-cm-long fine tungsten rod (0.5 mm in diameter) was connected to the encoder, serving as the encoder shaft. The edges of the white plastic cylinder limited the butterflies’ visual access to the sky to 110°. A small plastic tube (20 mm in length, 2 mm in diameter, with a 0.5 mm hole) was attached to the distal end of the rod, enabling the butterfly to be quickly and securely connected to the rod;
3. **a video recording system:** Two miniature video cameras (Waveshare Electronics, model H) were symmetrically positioned around the tungsten rod, with one recording the butterflies’ behaviour from above. The use of two cameras, as opposed to one, ensured a symmetrical visual environment within the flight simulator. A Raspberry Pi 4 single-board computer, housed in a grounded aluminum box along with a power bank, was used for real-time observation of red admiral flight behaviour and for storing raw video from each test for further analysis.

### Behavioural procedures

#### Indoor behavioural procedures

All flight simulator experiments under artificial light conditions were conducted in the nonmagnetic laboratory house (outside the main building of the station) with controlled temperature (21-22 °C) which was used in our previous magnetoreception studies on migratory birds (Bojarinova et al., 2020). The house comprised two rooms: the larger room (4 m × 4 m × 4 m) functioned as the primary experimental chamber. Its walls were plated with aluminium and grounded to effectively form a Faraday cage. A double-wrapped, three-dimensional Merritt four-coil system (‘magnetic coils’ hereafter) was positioned in this room to enable manipulation of the magnetic field during the experiments (Figure S1D). This Merritt coil system generates a magnetic field with over 99% homogeneity within a volume of approximately 110 × 110 × 110 cm, at the centre of which we placed the flight simulator. Each of the three axes of the coils was powered by a separate power supply BOP 50-4 M (Kepco Inc., USA), allowing precise control of the field’s intensity and direction. The parameters of the magnetic field were checked in the centre of the flight simulator using a FMV400 portable magnetometer (Meda Inc., USA) before tests. Professional LED strips (Arlight, Russia) were used as an artificial light source (395 nm UV LEDs + White LEDs (5000 K, CRI = 95-98), ∼ 8×10^18^ quanta s^−1^ m^−2^, see spectrum in Figure S1E). LED strips were mounted on an aluminum frame and covered with a layer of half-white photographic LEE filters to diffuse the LED light without altering its spectrum. They were powered by a Rigol DP711 programmable DC supply (Rigol Technologies, Inc., Beaverton, USA). The LI-1500 lightmeter (LI-COR Inc., USA) with a LI-190R terrestrial quantum sensor was used to measure the level of light intensity (in quanta) inside the flight simulator at the position of a tethered butterfly. We measured the spectrum of LEDs using the MK350N PREMIUM handheld spectrometer (UPRtek Inc., Taiwan). All electrical equipment used in the experiments, such as the power supply for LED light inside the main chamber and the power supplies for the magnetic coils, was grounded similarly to the behavioural set-up and housed in a smaller room adjacent to the laboratory with the behavioural setup. The level of radio-frequency noise, which could theoretically affect the magnetic sense of red admirals, was measured during both indoor and outdoor experiments using a calibrated active loop antenna FMZB 1512 (Schwarzbeck Mess-Elektronik, Germany) and a spectrum analyzer Rigol DSA815-TG (Rigol Technologies, USA).

#### Outdoor behavioural procedures

All flight simulator experiments under full-spectrum natural light conditions were conducted at an open experimental site where local landmarks (such as trees and buildings) were positioned far away so that they were invisible to the tested butterfly from within the simulator. Butterflies had access to the undisturbed natural magnetic field (intensity 50507 ± 48 nT (mean ± SD) and inclination 69.9° ± 0.6°). Red admirals were tested under clear skies (≤ 10–20% cloud cover) with direct visibility of the sun, in calm to light wind conditions (≤ 1–2 m/s), and at warm temperatures ranging from 19–25 °C. Temperature (inside the flight simulator) and wind/airflow speed were checked before each test using a custom-made Arduino-based weather station (see details here: https://magbbb.com/openscience/smartlab/) and a portable anemometer (CEM DT-618, Shenzhen Everbest, China), respectively.

#### Test preparations and common procedures for all experimental conditions

Before the tests, all butterflies caught on a given day were randomly divided into several groups according to the number of experimental conditions specific to the seasons and years (see details in subsequent sections). This approach allowed to minimize the effects of the progress of season and/or population-specific differences in migratory direction or flight motivation on results of tests. Prior to attachment to the tethering stalks, each red admiral was kept in a fridge for 10-15 min to immobilise it. A small (3 cm) section of tungsten rod (ferromagnetic free) with footplate attached to a small piece of paper (a tethering stalk here and after) was vertically glued to butterfly’s thorax. After this procedure, red admirals were fed and returned to a mesh cage, where they were kept for at least one more day to recover before being tested in the flight simulator. All flight tests took place between 10:00 and 17:00 hours local time (GMT+2) when the sun was visible from inside the simulator. Before experiments, mesh cages containing the butterflies were placed at the experimental site in direct sunlight for at least 30 minutes. This procedure allowed them to warm up and sense different orientation and navigation cues before testing. During testing, each red admiral was carefully removed from the mesh cage by grasping the stalk attached to its thorax. The stalk was then inserted into a small plastic tube attached to the rod in the flight simulator, as described in previous sections. Each test lasted 15 minutes and was included in the analysis only if 1) the butterfly flew for at least 5 minutes and 2) flapping was vigorous, with a large and symmetrical amplitude for both wings, indicating that the stalk was securely attached to the butterfly’s thorax and did not interfere with its flight behaviour (no formal criteria, the assessment of this was done by AP visually). To minimize handling bias, the first minute of each flight was excluded from the orientation analysis. The flight simulator was randomly rotated between trials to eliminate any potential directional biases associated with the apparatus.

### Experimental conditions

#### Experiments under full-spectrum natural light and artificial LED light during autumn migration 2023-2024 (indoors and outdoors)

The aim of these experiments was to determine whether migratory red admirals use geomagnetic field parameters as orientation cues. To investigate this, we conducted flight simulator experiments under varying light and magnetic conditions during August–September 2023–2024.

1. outdoors, with full access to the normal magnetic field (NMF), the sun and sky cues, full-spectrum natural light, autumn 2023-2024 (**Autumn Sun group**);
2. outdoors, the NMF, no access to the sun and the sky (overcast condition simulated with diffusor), full-spectrum natural light, autumn 2024 (**Autumn Overcast group**);
3. indoors, the NMF, artificial light (UV+white LEDs), autumn 2023-2024 (**Autumn NMF group**);
4. indoors, the changed magnetic field (CMF), artificial light (UV+ white LEDs), autumn 2023 (**Autumn CMF group**).

The CMF generated by the magnetic coils had a similar intensity as the NMF (50470 ± 180 nT and 50455 ± 145 nT, respectively), but the vertical component of the field was inverted (NMF inclination: 70.1 ± 0.25°, CMF inclination: −70.2 ± 0.13°). To simulate overcast conditions, where the butterflies could not view the sun or other sky cues (spectral composition or gradient of light intensity in the sky), the top of the simulator was covered with a layer of UV-transmissive diffusing paper (LEE diffusion gel filter). This setup reduced light intensity (∼2.4 × 10¹ quanta s ¹ m ² under overcast conditions versus ∼12.4 × 10¹ quanta s ¹ m ² under clear skies) without altering the spectrum of natural light (Figure S1E) which can be crucial for magnetoreception in butterflies (Guerra et al., 2014). Due to the absence of specialised tools to measure polarization of light inside this overcast setup, we cannot confirm whether red admirals could use polarized light as a cue during overcast experiments or exclude this possibility.

#### Experiments during spring migration 2023

To date, only one study based on visual observations has reported a northward flight direction in free-flying red admirals during spring migration, and no lab-controlled experiments have been conducted on this species in spring. To address this gap, we captured red admirals in May 2024 and randomly tested them under one of two experimental conditions:

1. outdoors, with full access to the normal magnetic field (NMF), the sun and sky cues (**Spring Sun group**);
2. indoors, the NMF, artificial UV + White LEDs light (**Spring NMF group**).

*Clock-shift experiments during autumn 2024 (outdoors)*.

To investigate the time-dependency of the sun compass in red admirals, we placed wild-caught butterflies in a windowless room with controlled temperature (21–22 °C) and an artificial light source, as described in Capture and Husbandry section. These butterflies were subjected to a 6-hour advanced clock-shift condition relative to local daylight time (lights on 6 hours after local sunrise and off 6 hours after local sunset; **Clock-shift group**). They were housed under conditions similar to those used for the control sun experiments mentioned earlier, including identical mesh cages, constant temperature, and controlled humidity, for a minimum of 7 days before the first test. The sun’s apparent azimuth movement across the sky averages approximately 15° per hour. Theoretically, if red admirals possess a time-dependent sun compass, their orientation would shift approximately 90° clockwise compared to the control group after a 6-hour forward clock shift. However, the azimuthal speed of the sun varies throughout the day, depending on the time and geographic location: actual 6-h azimuth movement of the sun at our experimental site varied between 92° and 104° (mean = 99°) based on data from the free software Stellarium. The flight simulator experiments for Clock shift group followed the same design as outdoor tests with Control group but started at least 1 hour after artificial sunrise time.

#### Experiments with the simulated sun during autumn migration 2024 (indoors)

To determine which cues hold a higher position in the hierarchy of compass systems in red admirals, we created a cue conflict between visual cues and the geomagnetic field. To achieve this, we slightly modified our flight simulator by mounting three green light-emitting diodes (530 nm peak, ∼ 3 × 10^17^ quanta s ¹ m ², see spectrum here (Romanova et al., 2023) on the inner wall at positions of 0°, 120°, and 240° relative to magnetic North (to create a symmetrical environment).

Previous studies on insects have shown that green LEDs can serve as a simulated sun cue. Insects, such as monarch butterflies and fruit flies, may either orient themselves towards the simulated sun or maintain arbitrary headings, when they consistently maintain a fixed but nonspecific direction relative to the position of the green cue (Giraldo et al., 2018; Franzke et al., 2020; Franzke et al., 2022; Beetz et al., 2023). The inner wall was covered with a black curtain to eliminate reflections from the green LEDs, which could otherwise create light artefacts. One group of red admirals was tested in the NMF with the green LED positioned at 0° (**N_GreenLED group**), while another group was tested with the LED at 120° (**SE_GreenLED group**). The background light intensity and spectrum were identical to those used in the indoor experiments.

### Data acquisition and statistical analysis

To study orientation of red admiral under different light and magnetic conditions, we used a video file as a primarily source of directional data. All video were analysed in DeepLabCut, an open-source software for markerless pose estimation (Mathis et al., 2018; Nath et al., 2019), to extract coordinates of body and head of a tested butterfly on each frame. Using these coordinates and custom-written Python scripts (see **Data availability** section for links), we calculated the mean flight direction (α) of the individual red admiral and individual r value (mean orientation vector length), which indicates the directedness of tested animal (its tendency to fly in the same direction; r value is between 0 and 1).

The classical Rayleigh test of uniformity (RT) was used to compare the group mean orientation against uniformity (Batschelet, 1981). Additionally, we were able to conduct second order statistics for this purpose - the nonparametric Moore’s modified Rayleigh test (MMRT; (Moore, 1980) which allows to weight the mean angles according to their r value. For both tests (RT and MMRT) low p-value (p < 0.05) indicates that the tested butterflies chose a preferred direction (modified version of R script from Massy et al., 2021). The nonparametric Mardia-Watson-Wheeler (MWW test) test was used to test the significance of the difference between orientation of red admirals from different experimental groups. We conducted maximum likelihood analysis for our samples using the CircMLE R package (Fitak and Johnsen, 2017) to specifically describe the patterns of orientation under different experimental conditions. Additionally, we used the bootstrap technique to identify whether oriented groups showed significantly more directed behaviours than non-significantly oriented groups. According to this method, a random sample of orientation directions (n angles) was drawn with replacement from the sample of orientation directions present in the significantly oriented group (Leberecht et al., 2022; Romanova et al., 2023). Based on these n orientation angles, the corresponding r-value was calculated, and this procedure was repeated 100,000 times. After that, the resulting 100,000 r-values are ranked in ascending order: the r-values at rank 2500 and 97500, at rank 500 and 99500 define the 95% and 99% limits for the actually observed r-value of the significantly oriented group, respectively. If the actual observed r-value of the disoriented group is outside these confidence intervals, the oriented group is significantly more directed than the disoriented group with a significance of p < 0.05 and p < 0.01, respectively. Bootstrap and maximum likelihood analyses, as well as MMRT, were conducted using R 4.3.2 (The R Core Team, 2013). The MWW test was performed in Oriana 4.02 (Kovach Computing Services, UK), while RT calculations and orientation visualisations for each group were carried out using a custom-written Python script.

## Results

### Orientation with access to the sun and the natural sky

When red admirals have full access to the natural sky and solar cues (such as sun position, polarized light, spectral contrast, and an intensity gradients) along with an undisturbed magnetic field, they typically exhibit the correct migratory direction in flight simulator experiments. The orientation of control red admirals with full access to natural cues was axially bimodal northwards and southwards during spring migration of 2024 (RT: α = 171±180°, r = 0.43, n = 25, 95% CI = 152°–190° and 332° - 10°, p = 0.01, Figure 1A; MMRT: Table S4A; CircMLE modelling results: Table S2A). During the autumn migration of 2023–2024, butterflies were oriented in a southeastern direction (RT: α = 169°, r = 0.59, n = 28, 95% CI = 144°–193°, p < 0.001, Figure 2A; MMRT: Table S4B; CircMLE modelling results: Table S2B). Similar results were observed in our previous study, where red admirals were tested under the same conditions at the same location during the autumn migrations of 2020–2021 (Pakhomov et al., 2023). In that study, butterflies exhibited a more southwestern direction (α = 198°); however, the difference in orientation between 2020–2021 and 2023–2024 was not statistically significant (MWW test: W = 0.659, p = 0.719) and mean vector strength was the similar between these groups (Kruskal-Wallis test: χ2 = 0.012, p = 0.91).

**Figure 1.**
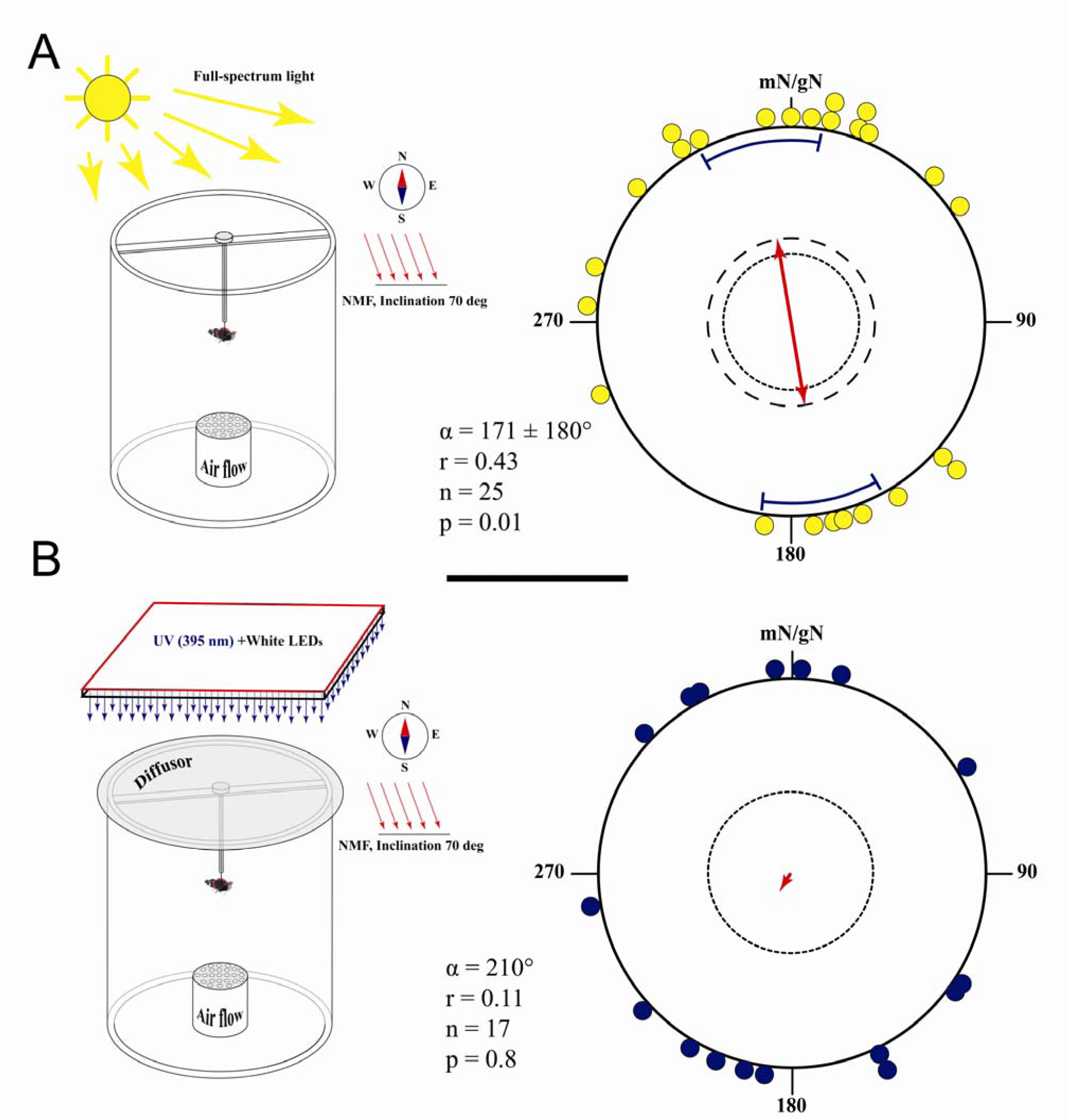
Orientation of red admirals under full-spectrum (A: outdoors, the natural sky and natural magnetic field) and artificial light (B: indoors, UV+ White LED and the natural magnetic field) conditions during spring migration 2023. **Left side:** schematic illustration of the experimental setup and experimental conditions. **Right side:** the flight directions of red admirals tested under full-spectrum (A) or LED (B) lights. Each dot at the circle periphery indicates the mean orientation of one individual butterfly. Dashed lines indicate the significance thresholds of the Rayleigh test for 5% and 1%, respectively. The mean direction of the admirals is shown as a red line, the curved lines indicate 95% confidence intervals. mN – magnetic North, gN – geographic North, NMF – the natural magnetic field.

**Figure 2.**
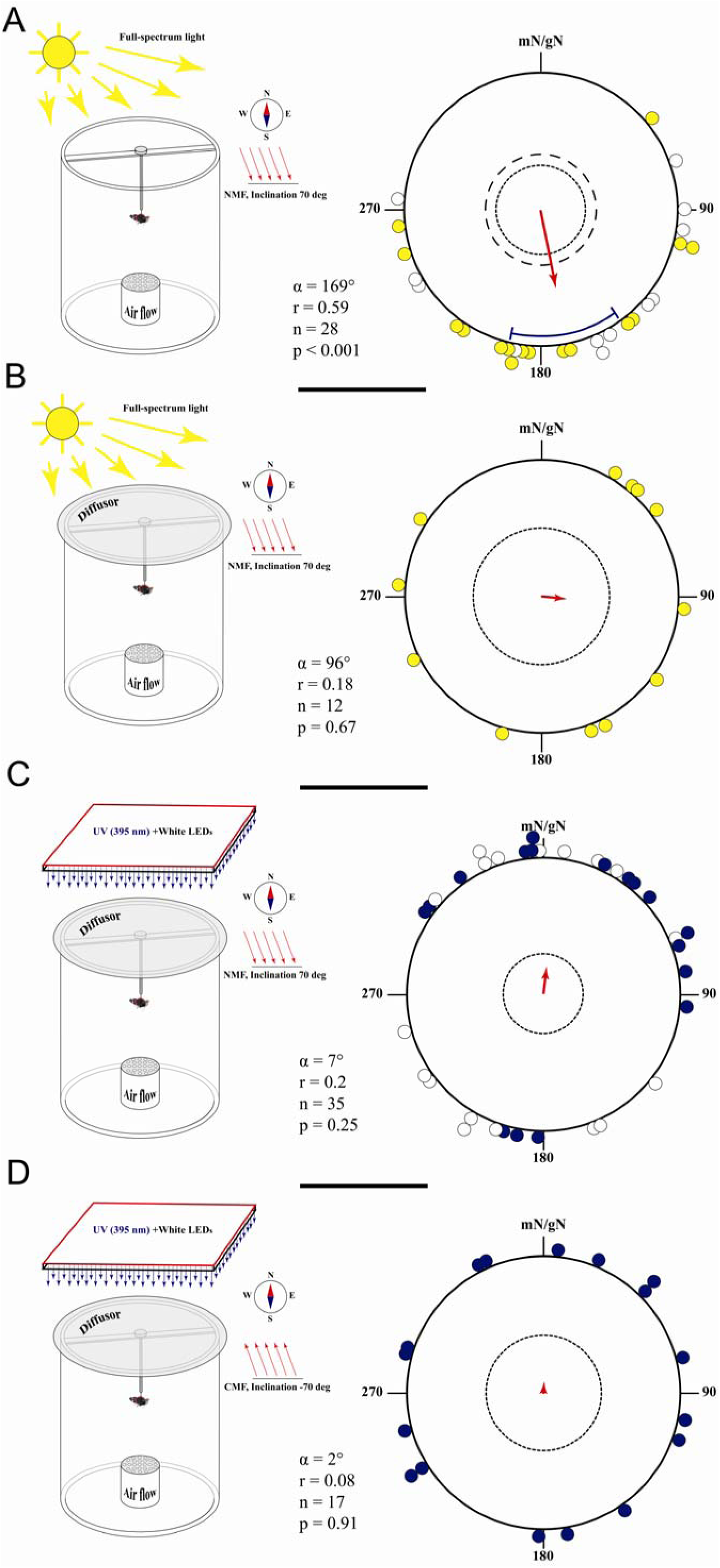
Orientation of red admirals under different light and magnetic conditions during autumn migration 2023-2024. A) outdoors, the natural sky, the sun and natural magnetic field (NMF), 2023: white dots, 2024: yellow dots; B) outdoors, simulated total overcast (diffusor) and natural magnetic field; C) indoors, UV+ White LED light and the natural magnetic field, 2023: purple dots, 2024: white dots d; D) indoors, UV+ White LED light and the changed magnetic field with inverted inclination of −70°. For details, see legend in Figure 1.

### Orientation without access to the sun and the natural sky

In the absence of any celestial cues, red admirals were unable to determine the appropriate migratory direction. In flight simulator experiments, they exhibited predominantly random flight activity, regardless of whether the light source was full-spectrum natural light (outdoors) or UV+White LED light (indoors).

### Outdoor experiment with the diffusor under natural light

If the upper part of the flight simulator placed outdoor under the sun was covered by a diffusor (overcast condition) which does not disturb the spectrum of sunlight but only decrease intensity of light and cut of all visual cues except the magnetic field, orientation of red admirals (autumn 2024) was not indistinguishable from the uniform distribution (RT: α = 98°, r = 0.18, n = 12, p = 0.67, Figure 2B; MMRT: Table S4C; CircMLE modelling results: Table S2C). The results of the bootstrap analysis indicate that the directions chosen by butterflies under overcast conditions were significantly more scattered compared to when their conspecifics had access to solar cues (Overcast_2024-Sun_2024: p < 0.01; 95% CI for r-value is 0.41 < r < 0.89; 99% CI for r-value is 0.35 < r < 0.96; Figure S6B).

#### Indoor experiment with the diffusor under artificial light

Similar results were observed in our indoor flight simulator experiments conducted in spring 2023: red admirals showed random flights under LEDs light in NMF (RT: α = 210°, r = 0.11, n = 17, p = 0.81, Figure 1B; MMRT: Table S4D; CircMLE modelling results: Table S2D). A more complex pattern emerged in the indoor experiments, where red admirals were tested under LED lighting and natural magnetic field conditions in autumn 2023-2024. According to the results of the classical Rayleigh test, red admirals were completely disoriented when exposed solely to magnetic cues under artificial light during the autumn migrations of 2023–2024 (α = 7°, r = 0.2, n = 35, p = 0.25; Figure 2C). Bootstrap analysis comparing all butterflies tested under natural sunlight versus those under artificial light revealed a significant difference (p < 0.01), with 95% confidence intervals for the r-value ranging from 0.45 to 0.73 and 99% confidence intervals from 0.42 to 0.80 (Figure S6A). However, a likelihood-based modeling approach in CircMLE (see Table S2E) and doubling the angles (RT_double_angles_: α = 26°, r = 0.29, n = 35, p = 0.051) suggest that the observed distribution might be close to bimodal. When analyzing the two years of experiments separately, red admirals reacted differently to this experimental condition. In 2023, they exhibited a tendency to orient in northeastern direction (RT: α = 28°, r = 0.4, n = 17, p = 0.065; MMRT: α = 33°, R* = 1.04, n = 17, 0.025< p < 0.05, Table S4E). In contrast, in 2024, red admirals were completely disoriented under the same conditions (RT: α = 292°, r = 0.14, n = 18, p = 0.695; MMRT: α = 346°, R* = 0.57, n = 18, 0.1 < p < 0.5; Table S4E), as well as their conspecifics tested under identical lighting but with a reversed magnetic inclination (−70° CMF) during the autumn migration of 2023 (RT: α = 2°, r = 0.08, n = 17, p = 0.91; Figure 2D; MMRT: Table S4F; CircMLE modelling results: Table S2F). Bootstrap results for the 2023 Sun_–70° CMF condition confirmed the loss of orientation (p < 0.01), with the 95% CI for the r-value ranging from 0.34 to 0.78 and the 99% CI from 0.3 to 0.88 (Figure S6C).

#### Indoor experiments with the diffusor and green LED

Adding visual cues, such as a green LED simulating the sun, improved the red admirals’ orientation ability in the indoor experimental setup. Butterflies in the N_GreenLED group did not fly directly toward the green LED (95% CI does not include the diode’s position; RT: α = 306°, r = 0.46, n = 18, p = 0.022, 95% CI = 267° - 350°, Figure 4A; MMRT: Table S4G; CircMLE modelling results: Table S2G). Instead, most butterflies kept the LED positioned on the right side of their visual field. When the position of the bright green LED was shifted 120° clockwise (from magnetic N to magnetic SE) and another group of admirals (SE_GreenLED group) was tested, the butterflies exhibited a similar directional pattern (RT: α = 60°, r = 0.5, n = 17, p = 0.01, 95% CI = 15° - 92°, Figure 4A; MMRT: Table S4H; CircMLE modelling results: Table S2H) and their orientation was different from that of N_GreenLED group (95 and 99% CIs do not overlap; but MWW test: W = 5.361, p = 0.07).

**Figure 3.**
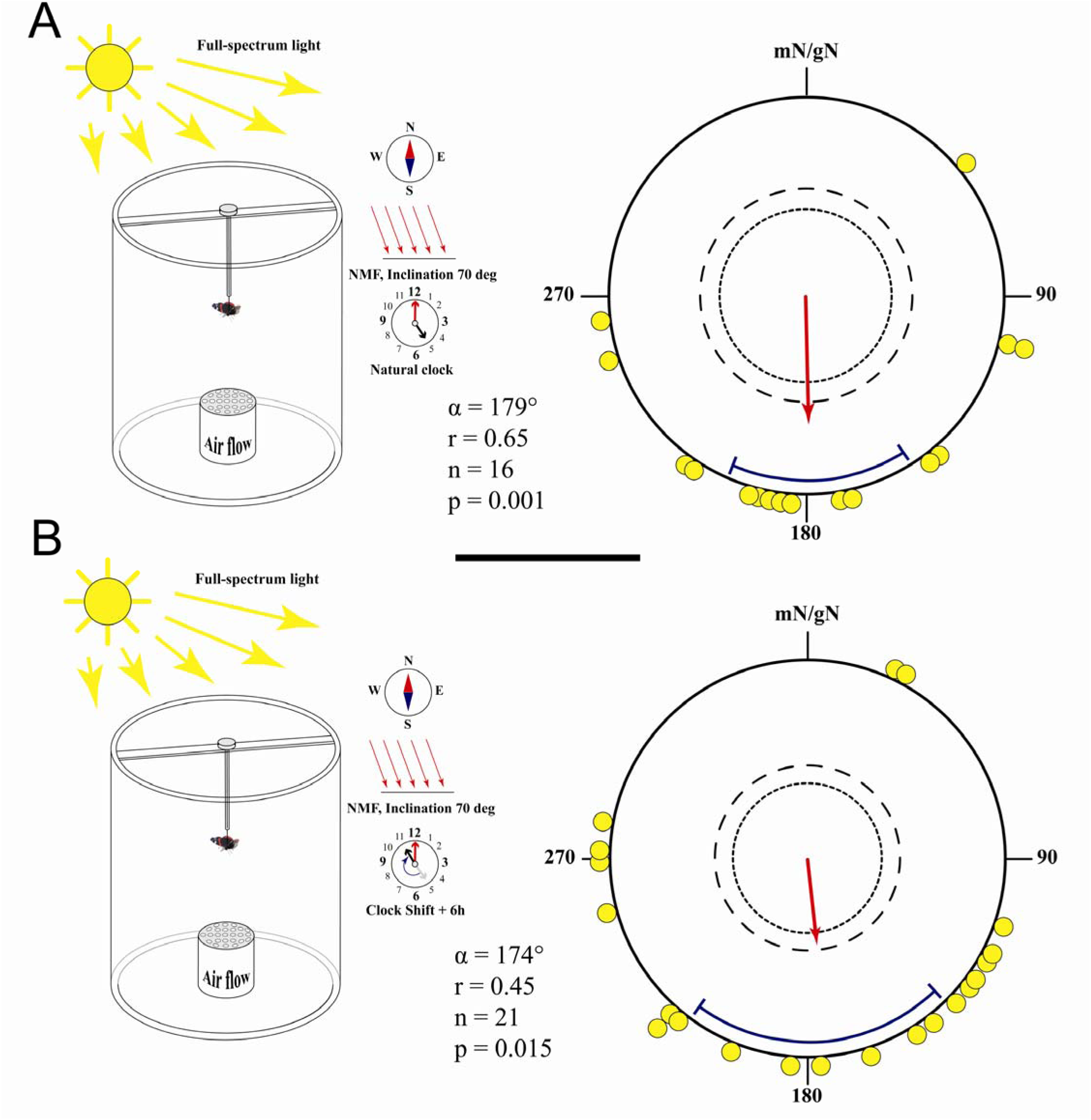
Orientation of red admirals after 6 hours forward clock shift during autumn migration 2024. A) control group: outdoors, the natural sky, the sun and natural magnetic field (NMF); B) clock shift group: outdoors, the natural sky, the sun and natural magnetic field (NMF). For details, see legend in Figure 1.

**Figure 4.**
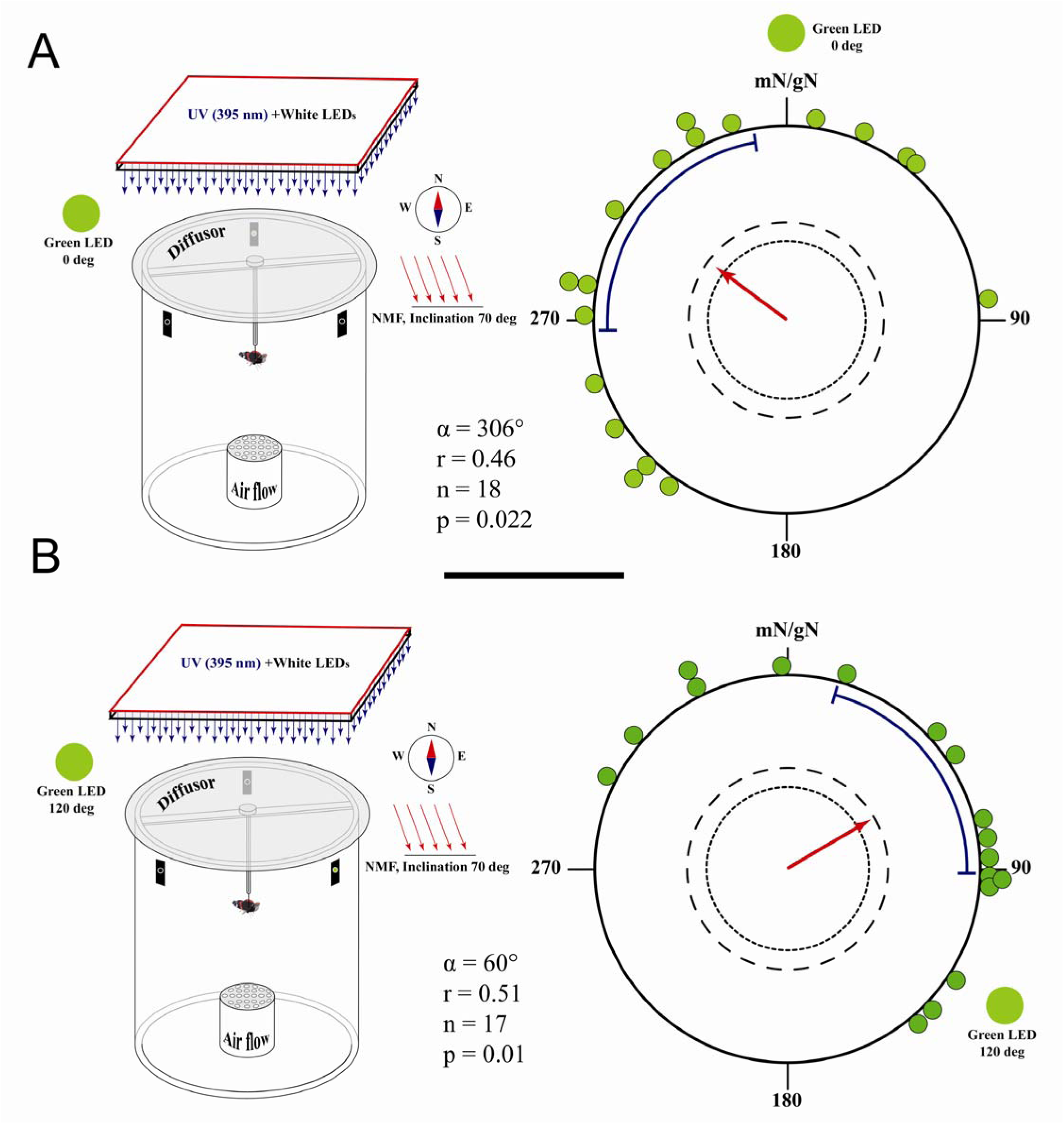
Orientation of red admirals under artificial light condition with a green LED as a simulated sun during autumn migration 2024. A) indoors, indoors, UV+ White LED light and the natural magnetic field, green LED is in 0° (mN) position; B) indoors, indoors, UV+ White LED light and the natural magnetic field, green LED is in 120° (mSE) position. For details, see legend in Figure 1.

### Orientation with access to the sun and the natural sky after 6-h clock shift

Typically, animals that rely on solar cues for orientation compensate for the sun’s azimuth changes throughout the day and, consequently, respond to clock-shift procedures. However, after being kept for at least a week under a 6-hour forward-shifted artificial photoperiod, all red admirals tested in the flight simulator during autumn 2024 exhibited a southward mean direction (RT: α = 174°, r = 0.45, n = 21, p = 0.015, 95% CI = 136° - 216°, Figure 3B; MMRT: Table S4K; CircMLE modelling results: Table S2K). This direction was similar to the mean direction of the non-clock-shifted butterflies (RT: α = 179°, r = 0.65, n = 16, p = 0.001, 95% CI = 148° - 204°, Figure 3A; MMRT: Table S4J; CircMLE modelling results: Table S2J). There is no difference in orientation between clock-shifted and control groups (MWW test: W = 3.56, p = 0.17) and mean vector strength was the similar between these groups (Kruskal-Wallis test: χ2 = 3.96, p = 0.15).

## Discussion

### Migratory direction of red admirals tested under natural condition

In the wild, red admirals exhibit a broad directional range from east to southwest during autumn migration across different parts of Europe, according to studies tracking the vanishing directions of freely flying butterflies (Williams, 1951; Roer, 1991; Benvenuti et al., 1994; Benvenuti et al., 1996; Leverton, 2000; Hansen, 2001; Brattström et al., 2008; see Figure 1 in Pakhomov et al., 2023 for a more detailed map). Little is known about their spring migration direction; however, one study based on visual observations in the wild found that red admirals tend to fly northward in southern Europe during spring (Benvenuti et al., 1996). Under laboratory-controlled conditions using flight simulators, our previous orientation experiments conducted on the Courish Spit confirmed a predominant south-westward orientation of red admirals during the autumn migration period (Pakhomov et al., 2023). In present study, red admiral tested in the flight simulator under sunny condition in August and September preferred a more south-southeast direction. During spring migration, the orientation of red admirals exposed to the natural magnetic field and a sunny, clear sky exhibited an axially bimodal pattern, which was unexpected. It is possible that our field site hosts a mix of migratory individuals flying north and local individuals flying in the opposite direction. According to the results of long-term monitoring using the two large traps described in the Methods section, there is a seasonal difference in the proportion of butterflies captured by the NE and SW traps. The NE trap, which is open to the south and primarily captures butterflies flying northward, and the SW trap, open to the north and capturing southward-flying butterflies, show differing catch ratios between spring and autumn migrations. In spring, similar numbers of butterflies are captured by both traps (2000–2022 data: see Figure 4 in Pakhomov et al., 2023), indicating bidirectional movements of free-flying butterflies at the study site during this migration season. However, in 2023-2024, approximately 75% of individuals were caught in the NE trap and only about 25% in the SW trap (Figure S1), suggesting a predominant northward movement during spring migration. In autumn, most red admirals are usually caught in the SW trap (Figure 4 in Pakhomov et al., 2023; Figure S1), indicating that they predominantly fly in the expected migratory southward direction, a pattern that is also confirmed by our flight simulator experiments. Bimodal orientation is not an uncommon phenomenon in animal migration and has been documented in both free-flying and laboratory-tested species (Åkesson et al., 1996; Schneider et al., 2023; Voigt et al., 2023). Despite the more complex flight behaviour during spring migration, this study provides the first demonstration of migratory orientation in red admirals tested under lab-controlled conditions during both migratory seasons. Orientation behaviour in spring appears to be more complex (axial north-south) compared to unimodal oriented behaviour demonstrated during autumn migration.

### Visual cues vs Magnetic field

Results of all our experiments under full-spectrum natural and artificial LED light conditions indicate that migratory red admirals mostly rely on visual solar cues when they choose and maintain seasonal directions. With the full access to the clear sky cues which contain a direct view of the sun, polarized light, spectral contrast, and light intensity gradients (Franzke et al., 2020; Franzke et al., 2022), butterflies prefer southward direction in autumn (Figures 2A, 3A) or show bimodal distribution with predominance of the northward group in spring (Figure 1A). When the view of the clear sky is restricted — either by covering the flight simulator with a diffuser in outdoor experiments (simulating overcast conditions) or by using an artificial light source in indoor tests — so that the geomagnetic field becomes the only available directional cue, this manipulation does not result in a dramatic decrease in the number of active butterflies (Figure S3). However, red admirals either become disoriented (Figure 1B, 2B, C, D) or exhibit an inappropriate migratory direction (Figure S4, E:2023). Analysis of long-term phenology data, collected from migratory animal traps along the eastern Baltic coast between 1983 and 2022 (Pakhomov et al., 2023), indicates that the abundance of red admirals strongly correlates with clear sky conditions. Of the 20,672 individuals recorded, 44.2% were captured under clear skies (0–25% cloud cover), 47.8% under partially cloudy conditions (25–50% cloud cover), and only 8% when cloud cover exceeded 75%, with just 1% recorded under total overcast conditions. Visual observation studies conducted in various parts of Europe further support this trend, indicating that red admiral migration typically occurs on warm, clear, sunny days with favorable winds (Mikkola, 2003; Brattström et al., 2008). In contrast, unfavorable weather conditions, such as overcast skies and/or strong winds, tend to suppress their migratory activity. This contrasts with the migratory behaviour of North American monarch butterflies, whose flight motivation and bearings are not significantly affected by overcast conditions (Schmidt-Koenig, 1979). Adding additional visual cues (green LEDs as a simulated sun) in our indoor setup restored their directional flight (Figure 4) compared to the non-visual cue conditions (Figures 1A and 2C). As mentioned in the Methods section, under such conditions, insects usually orient towards the simulated sun or maintain a fixed but nonspecific direction relative to the position of the cue (Byrne et al., 2003; Giraldo et al., 2018; Franzke et al., 2022; Beetz et al., 2023). Red admirals, on other hand, mostly tried to keep the green LED on the right side of their visual field while flying inside the flight simulator. All of this leads to the conclusion that European red admirals do not steer migratory flight behaviour by sensing the Earth’s magnetic field, and restricting solar cues disrupts their directional preference.

### Compass system of the European Red Admiral: a working hypothesis

In contrast to the more complex navigational system of the iconic monarch butterflies, which relies on the time-compensated sun compass as the primary system and the light-dependent inclination magnetic compass as a backup, red admirals appear to use a different orientation strategy. Below, we present the first working hypothesis based on the current study results:

1. Sun compass

Red admirals use the sun compass as the primary orientation mechanism to select and maintain seasonal migratory direction. As shown in our clock-shift experiments, the sun compass of red admirals is not linked to their internal clocks, similar to painted lady butterflies (Nesbit et al., 2009). Instead, they likely use the sun as a simple heading indicator (Guilford and Taylor, 2014), either flying toward it (free-flying red admirals at Pape station; Pakhomov et al., 2023) or maintaining a fixed angle relative to it before and after noon, without compensating for its movement throughout the day. This feature contrasts their sun orientation with that of other well-studied diurnal migrants, such as monarch butterflies, hoverflies, some neotropical butterflies, and homing pigeons (Schmidt-Koenig, 1958; Oliveira et al., 1998; Mouritsen and Frost, 2002; Massy et al., 2021). Another possible explanation for these contradictory results, as proposed in the painted lady study mentioned above — and which may also apply to red admirals — is that the clock-shifting experimental treatment did not successfully reset their internal clock, or they may have recalibrated their internal clock just before the tests in the flight simulator.

1. Magnetic compass

Red admirals appear not to possess a magnetic sense or a functional inclination magnetic compass, at least at the population or group level, as they were disoriented in both natural (indoors and outdoors) and inverted magnetic fields in the absence of visual cues. Another European migratory butterflies, painted lady, showed similar results under overcast condition: they were not able to determine appropriate migratory direction when only the geomagnetic field was available (Nesbit et al., 2009). Therefore, these two European migratory butterflies share a common feature that distinguishes them from migratory monarchs: they are not able to extract information on migratory direction from the geomagnetic field. The magnetic compass of monarch butterflies is sensitive to specific light parameters, with the spectral range between 380 and 420 nm being critical for magnetic orientation (Guerra et al., 2014; Kendzel et al., 2023). Previous studies that reported an absence of magnetic orientation in monarchs (Mouritsen & Frost, 2002; Stalleicken et al., 2005) have been criticised for not providing appropriate light conditions (Guerra et al., 2014). In our study, we used a diffuser that does not alter the light spectrum in our outdoor experiments conducted under full-spectrum natural light, and we supplemented the white light source in our indoor experiments with UV LEDs (395 nm). However, even under these experimental conditions, red admirals remained disoriented, which may indicate the absence of a magnetic sense in this species.

1. Why do red admirals’ orientation mechanisms differ from monarch butterfly?

Both migratory monarch butterflies in North America and Bogong moths in Australia can be considered specialists: their multigenerational life cycles depend on critical factors such as specific non-breeding sites (e.g., a small region of oyamel fir forests for monarch’s hibernation or alpine caves for moths’ aestivation). This reliance makes them highly vulnerable to rapid environmental changes caused by human activity (Thogmartin et al., 2017; Tenger-Trolander et al., 2019; Green et al., 2021; Parlin et al., 2022) and may have driven the evolution of a more complex navigation system incorporating at least two compass mechanisms (magnetic and astronomical). By contrast, red admirals rely on a widely available host plant — nettle — and do not migrate to a specific overwintering region in Mediterranean region (Stefanescu, 2001; Cuadrado, 2017). Given this ecological flexibility, they exhibit regular migrations with no significant decline in numbers (Pakhomov et al., 2023) and it is not surprising that they do not employ multiple complex compass mechanisms to select and maintain their seasonal migratory direction but instead rely on a simpler orientation mechanism.

### Conclusion

There are hundreds, if not thousands, of migratory butterfly and moth species around the world, yet most of them likely remain understudied. Only a small fraction of these species has been documented in the literature and the available studies are clearly biased towards monarchs (Chowdhury et al., 2021). However, even a small number of studies on migratory Lepidoptera have revealed that these migratory insects employ a variety of strategies for orientation. For example, North American monarch butterflies use the bird-like light-dependent inclination compass, European red admirals and painted lady butterflies seem to lack a magnetic sense, and Australian Bogong moths demonstrate a complex interaction between visual landmarks and the magnetic compass. More researches are needed on different species with varying migration strategies to deepen our understanding of their migrations and how human activities may negatively impact them.

## Acknowledgments

We are grateful to Roman Cherbunin for his help in measurements of radio-frequency noise and to Fyodor Cellarius for his help with video and statistical analysis (Python scripts).

## Authors’ contributions

A.P. and D.K. designed flight simulator research; A.P. performed experiments in a flight simulator, collected and analysed the data; N.S. and A.S. collected butterflies and analysed phenology data; A.P., N.S and D.K. wrote the first draft of the manuscript. All authors commented on the manuscript and gave final approval for publication.

## Data availability statement

All data (raw data of experiments, results of additional statistical analysis, etc) and Python/R scripts are provided in electronic supplementary material (https://doi.org/10.5281/zenodo.15316518). All components of the flight simulator, video analysis scripts, 3D printing/laser cutting files can be found here: https://magbbb.com/openscience/smartlab/flight-simulator/.

