## Supplementary materials for "Not All Butterflies Are Monarchs: Compass Systems in the Red Admiral (*Vanessa atalanta*), a European Diurnal Migrant"

**Figure S1. Experimental setups and the large ‘Rybachy-like’ traps for butterflies’ catching. A)** Directional preferences of movements of red admirals (up: number of individuals, middle: photo of traps from a drone, down: proportions) captured in two large traps in 2023-2024.

| **A** | 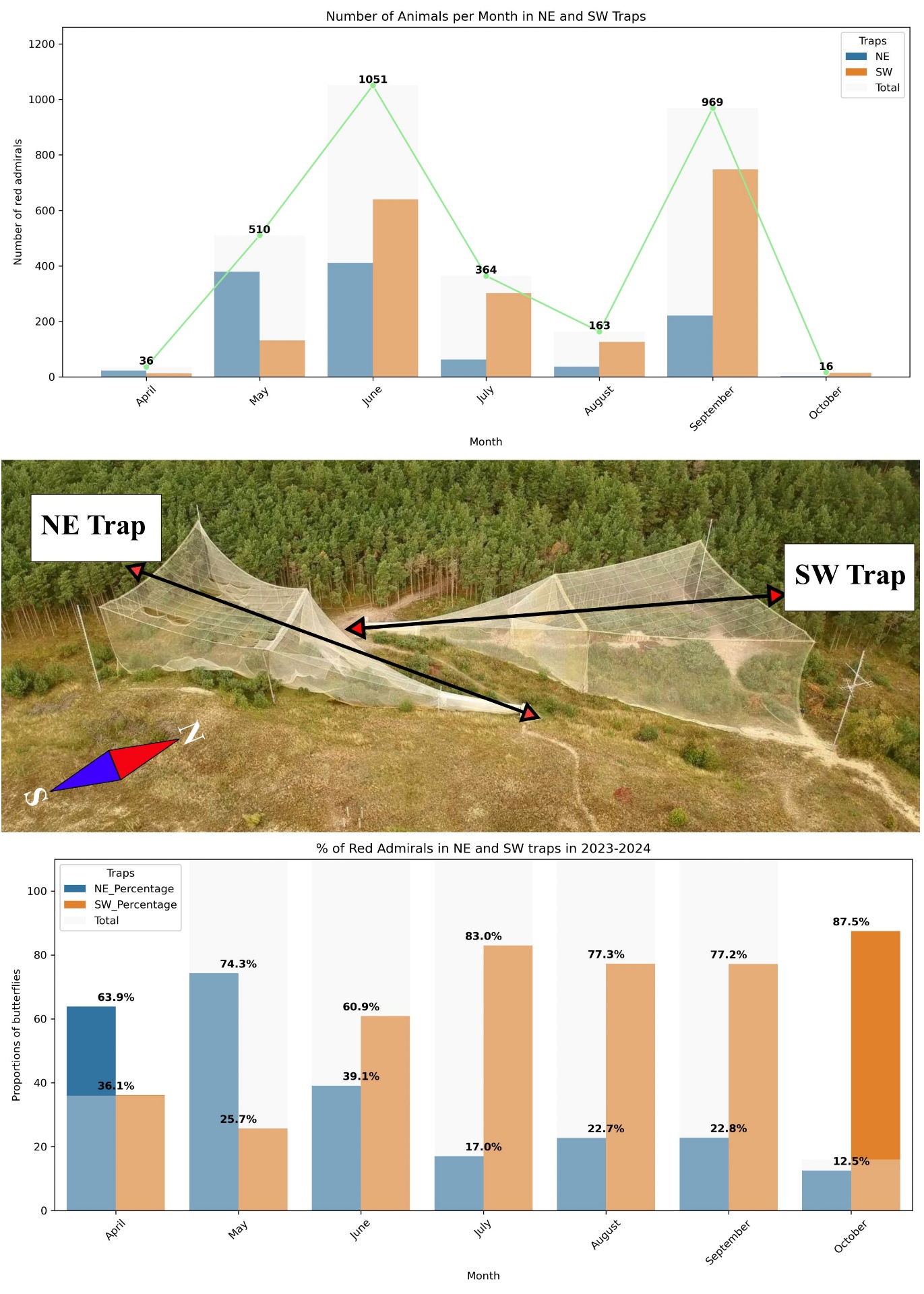 |
| --- | --- |

**Figure S1 (continue): B-C) A photo (B) and scheme (C) of the modified version of Mouritsen-Frost flight simulator:** 1 - an optical encoder, 2 - a miniature camera, 3 - Raspberry Pi 4, 5 - a PWN computer fan, 6 – a laptop; **D) Experimental setup for indoor experiments. E) Spectrometric measurements of UV+White LEDs light used in indoor experiments: blue** (in the laboratory chamber) and full-spectrum natural light outdoors (**red**: under sun condition, **orange**: under overcast condition)**; F) Average amplitude of the magnetic field noise measured at the outdoor experimental site as a function of the central position of the 10 kHz detection window**. 1 – near the box with minicomputer Raspberry PI (2 m away from the flight simulator), 2 - inside the flight simulator, the place where red admiral was attached to a vertical encoder shaft; all electronic equipment was turn on, 3 - inside the flight simulator, the place where red admiral was attached to a vertical encoder shaft; all electronic equipment was turn of.

| **B** | **C** | **D** |
| --- | --- | --- |
| 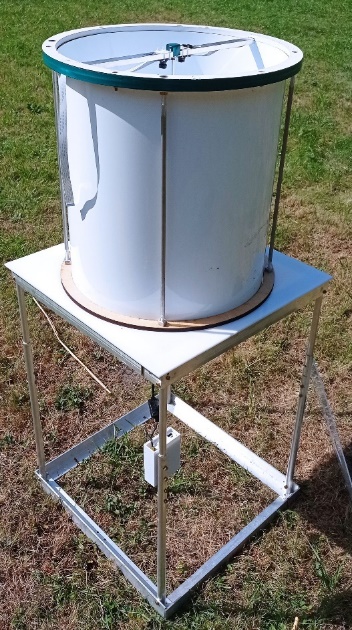 | 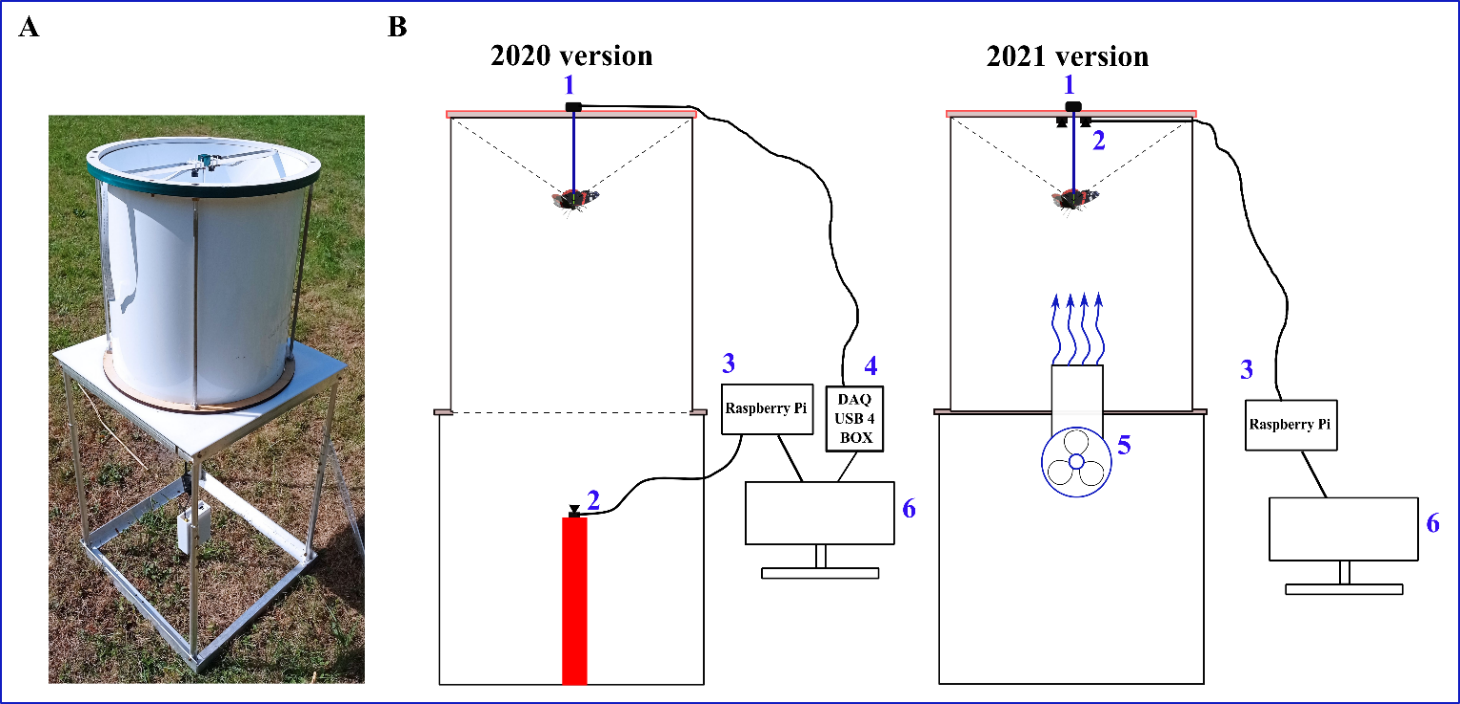 | 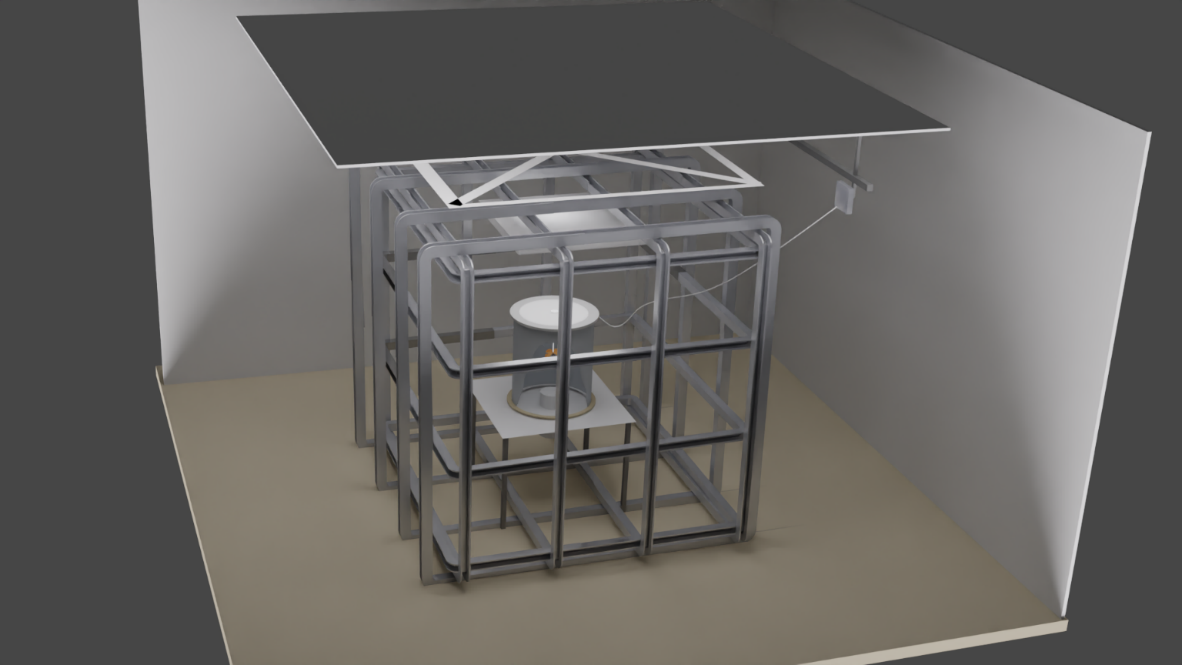 |
| **E** | | **F** |
| 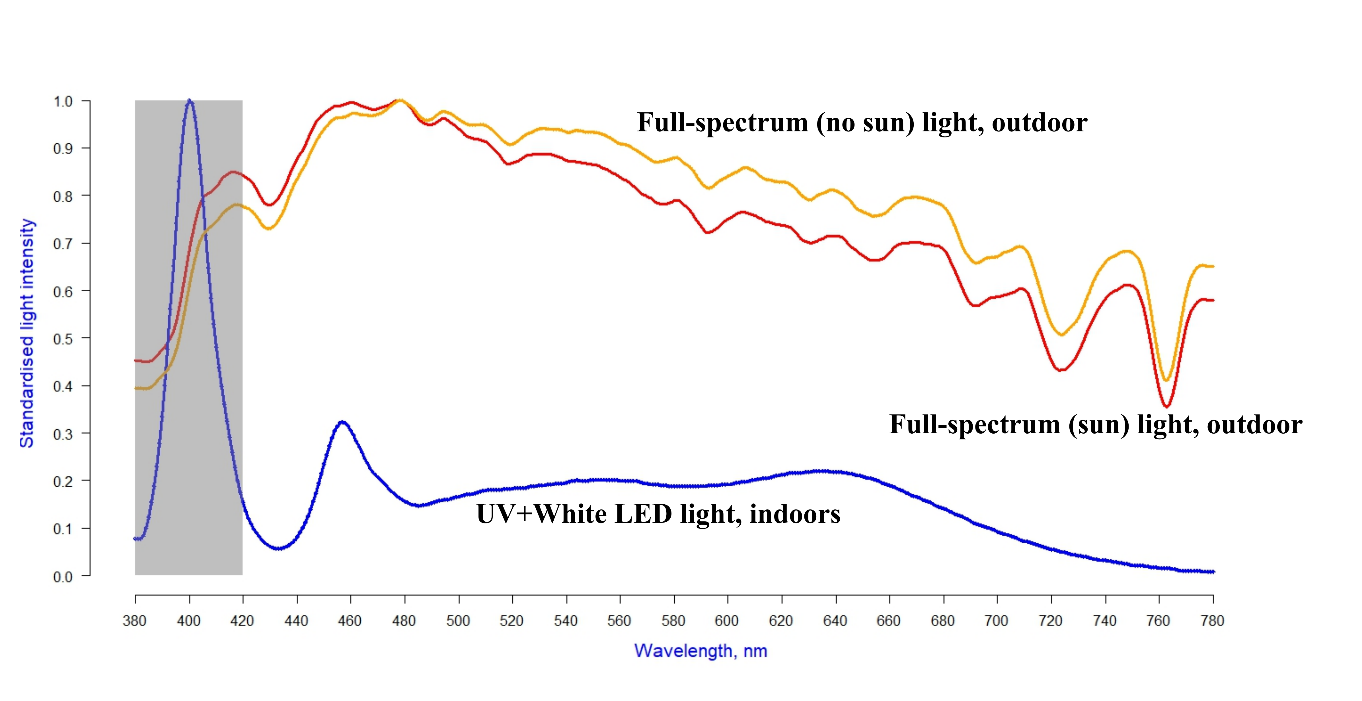 | | 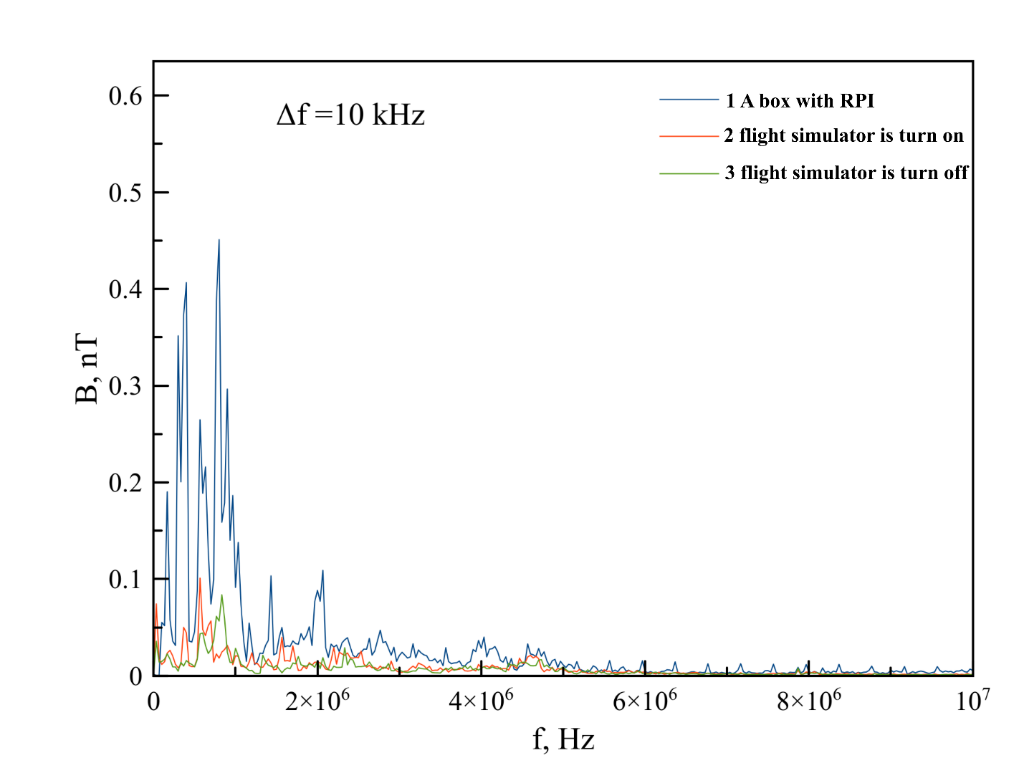 |

| **Figure S2.** Average amplitude of the magnetic field noise measured at the laboratory chamber as a function of the central position of the 10 kHz detection window. A – all electronic equipment OFF, B – all electronic equipment ON, ∆ f = 10 kHz | | |
| --- | --- | --- |
| **A** | 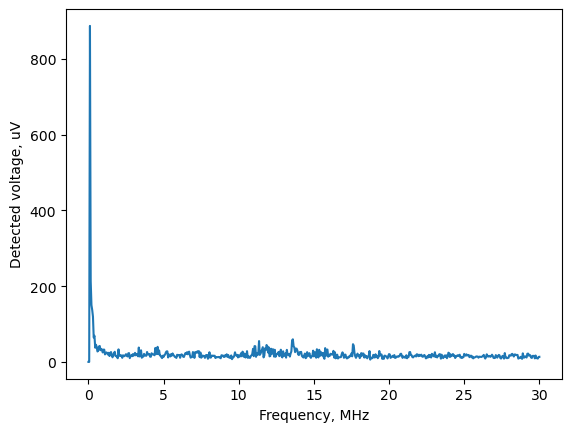 | 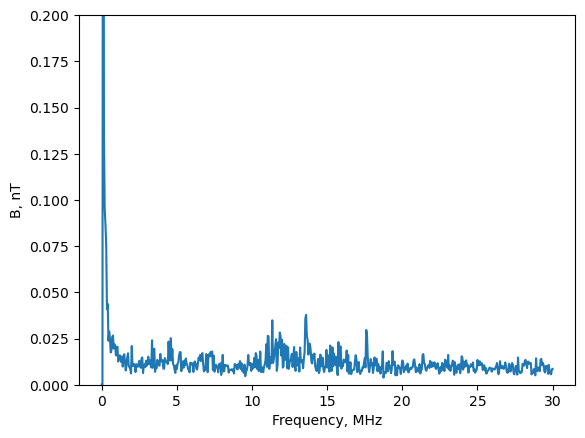 |
| **B** | 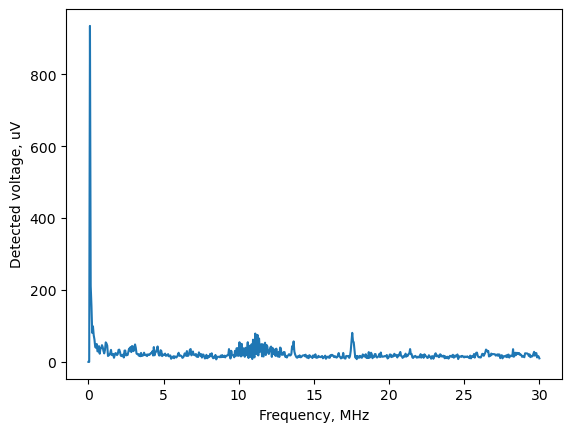 | 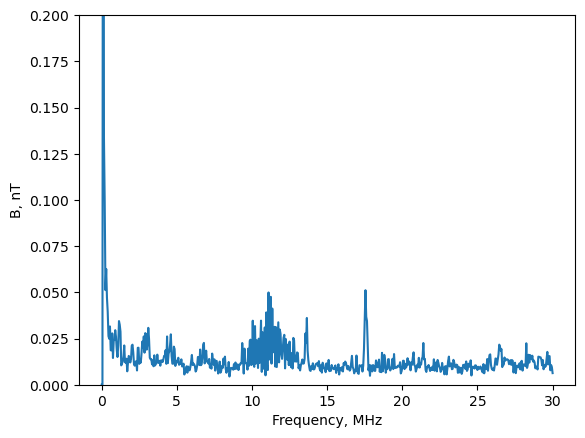 |

**Table S1. Raw data of all orientation tests.**

| **Year** | **Season** | **n all tested admirals** | **№ of active admirals** | **Magnetic condition** | **Light source** | **Mean direction, α** | **Length of mean vector, r** |
| --- | --- | --- | --- | --- | --- | --- | --- |
| **2024** | **Spring** | **57** | 1 | **Natural magnetic field** | **Full-spectrum light (outdoors, sun)** | 287 | 0.55 |
|  |  |  | 2 |  |  | 275 | 0.39 |
|  |  |  | 3 |  |  | 310 | 0.2 |
|  |  |  | 4 |  |  | 327 | 0.71 |
|  |  |  | 5 |  |  | 328 | 0.68 |
|  |  |  | 6 |  |  | 12 | 0.2 |
|  |  |  | 7 |  |  | 334 | 0.84 |
|  |  |  | 8 |  |  | 161 | 0.49 |
|  |  |  | 9 |  |  | 173 | 0.4 |
|  |  |  | 10 |  |  | 167 | 0.93 |
|  |  |  | 11 |  |  | 131 | 0.24 |
|  |  |  | 12 |  |  | 249 | 0.46 |
|  |  |  | 13 |  |  | 0 | 0.38 |
|  |  |  | 14 |  |  | 12 | 0.47 |
|  |  |  | 15 |  |  | 148 | 0.45 |
|  |  |  | 16 |  |  | 166 | 0.93 |
|  |  |  | 17 |  |  | 131 | 0.2 |
|  |  |  | 18 |  |  | 19 | 0.2 |
|  |  |  | 19 |  |  | 352 | 0.41 |
|  |  |  | 20 |  |  | 5 | 0.43 |
|  |  |  | 21 |  |  | 187 | 1 |
|  |  |  | 22 |  |  | 44 | 0.46 |
|  |  |  | 23 |  |  | 20 | 0.24 |
|  |  |  | 24 |  |  | 22 | 0.54 |
|  |  |  | 25 |  |  | 56 | 0.4 |
| **2024** | **Spring** | **28** | 1 | **Natural magnetic field** | **Full-spectrum light (indoors, UV + White LEDs)** | x | 0.67 |
|  |  |  | 2 |  |  | 125 | 1 |
|  |  |  | 3 |  |  | 356 | 1 |
|  |  |  | 4 |  |  | 124 | 0.68 |
|  |  |  | 5 |  |  | 203 | 0.85 |
|  |  |  | 6 |  |  | 313 | 0.13 |
|  |  |  | 7 |  |  | 193 | 0.82 |
|  |  |  | 8 |  |  | 187 | 0.46 |
|  |  |  | 9 |  |  | 155 | 1 |
|  |  |  | 10 |  |  | 260 | 0.2 |
|  |  |  | 1 |  |  | 209 | 1 |
|  |  |  | 12 |  |  | 333 | 0.3 |
|  |  |  | 13 |  |  | 153 | 0.8 |
|  |  |  | 14 |  |  | 15 | 0.88 |
|  |  |  | 15 |  |  | 331 | 0.53 |
|  |  |  | 16 |  |  | 4 | 0.4 |
|  |  |  | 17 |  |  | 226 | 0.68 |
| **2024** | **Autumn** | **28** | 1 | **Natural magnetic field** | **Full-spectrum light (outdoors, sun)** | 104 | 0.6 |
|  |  |  | 2 |  |  | 197 | 0.26 |
|  |  |  | 3 |  |  | 186 | 0.21 |
|  |  |  | 4 |  |  | 216 | 0.4 |
|  |  |  | 5 |  |  | 169 | 0.6 |
|  |  |  | 6 |  |  | 213 | 0.14 |
|  |  |  | 7 |  |  | 194 | 0.5 |
|  |  |  | 8 |  |  | 104 | 0.41 |
|  |  |  | 9 |  |  | 252 | 0.24 |
|  |  |  | 10 |  |  | 262 | 0.42 |
|  |  |  | 11 |  |  | 189 | 0.21 |
|  |  |  | 12 |  |  | 141 | 0.6 |
|  |  |  | 13 |  |  | 172 | 0.7 |
|  |  |  | 14 |  |  | 144 | 0.43 |
|  |  |  | 15 |  |  | 188 | 0.67 |
|  |  |  | 16 |  |  | 49 | 0.73 |
| **2024** | **Autumn** | **42** | 1 | **Natural magnetic field** | **Full-spectrum light (outdoors, sun), Clock Shift +6h** | 126 | 0.45 |
|  |  |  | 2 |  |  | 186 | 0.48 |
|  |  |  | 3 |  |  | 201 | 0.62 |
|  |  |  | 4 |  |  | 27 | 0.52 |
|  |  |  | 5 |  |  | 129 | 0.75 |
|  |  |  | 6 |  |  | 143 | 0.24 |
|  |  |  | 7 |  |  | 134 | 0.32 |
|  |  |  | 8 |  |  | 117 | 0.58 |
|  |  |  | 9 |  |  | 25 | 0.47 |
|  |  |  | 10 |  |  | 109 | 0.45 |
|  |  |  | 11 |  |  | 254 | 0.48 |
|  |  |  | 12 |  |  | 220 | 0.95 |
|  |  |  | 13 |  |  | 280 | 0.93 |
|  |  |  | 14 |  |  | 271 | 0.86 |
|  |  |  | 15 |  |  | 269 | 0.46 |
|  |  |  | 16 |  |  | 121 | 0.23 |
|  |  |  | 17 |  |  | 219 | 0.58 |
|  |  |  | 18 |  |  | 221 | 0.25 |
|  |  |  | 19 |  |  | 177 | 0.76 |
|  |  |  | 20 |  |  | 148 | 0.62 |
|  |  |  | 21 |  |  | 162 | 0.43 |
| **2024** | **Autumn** | **32** | **1** | **Natural magnetic field** | **Full-spectrum light (indoors, UV + White LEDs)/ Green LED as a simulated sun (position 0 deg)** | 238 | 0.28 |
|  |  |  | 2 |  |  | 223 | 0.34 |
|  |  |  | 3 |  |  | 8 | 0.17 |
|  |  |  | 4 |  |  | 37 | 0.86 |
|  |  |  | 5 |  |  | 253 | 0.92 |
|  |  |  | 6 |  |  | 321 | 0.92 |
|  |  |  | 7 |  |  | 345 | 0.18 |
|  |  |  | 8 |  |  | 271 | 0.17 |
|  |  |  | 9 |  |  | 21 | 0.75 |
|  |  |  | 10 |  |  | 280 | 0.25 |
|  |  |  | 11 |  |  | 85 | 0.28 |
|  |  |  | 12 |  |  | 216 | 0.9 |
|  |  |  | 13 |  |  | 332 | 0.63 |
|  |  |  | 14 |  |  | 279 | 0.28 |
|  |  |  | 15 |  |  | 334 | 0.76 |
|  |  |  | 16 |  |  | 302 | 0.63 |
|  |  |  | 17 |  |  | 225 | 0.75 |
|  |  |  | 18 |  |  | 40 | 0.47 |
| **2024** | **Autumn** | **39** | 1 | **Natural magnetic field** | **Full-spectrum light (indoors, UV + White LEDs)/ Green LED as a simulated sun (position 120 deg)** | 47 | 0.16 |
|  |  |  | 2 |  |  | 17 | 0.33 |
|  |  |  | 3 |  |  | 122 | 0.51 |
|  |  |  | 4 |  |  | 332 | 0.74 |
|  |  |  | 5 |  |  | 334 | 0.72 |
|  |  |  | 6 |  |  | 140 | 0.5 |
|  |  |  | 7 |  |  | 75 | 0.56 |
|  |  |  | 8 |  |  | 93 | 0.3 |
|  |  |  | 9 |  |  | 92 | 0.1 |
|  |  |  | 10 |  |  | 358 | 0.36 |
|  |  |  | 11 |  |  | 95 | 0.78 |
|  |  |  | 12 |  |  | 296 | 0.87 |
|  |  |  | 13 |  |  | 82 | 0.74 |
|  |  |  | 14 |  |  | 55 | 0.25 |
|  |  |  | 15 |  |  | 135 | 0.36 |
|  |  |  | 16 |  |  | 88 | 0.63 |
|  |  |  | 17 |  |  | 312 | 0.57 |
| **2023-2024** | **Autumn** | **48** | 1 | **Natural magnetic field** | **Full-spectrum light (outdoors, sun)** | 240 | 0.44 |
|  |  |  | 2 |  |  | 128 | 0.98 |
|  |  |  | 3 |  |  | 71 | 0.22 |
|  |  |  | 4 |  |  | 150 | 0.28 |
|  |  |  | 5 |  |  | 156 | 0.27 |
|  |  |  | 6 |  |  | 157 | 0.45 |
|  |  |  | 7 |  |  | 239 | 0.44 |
|  |  |  | 8 |  |  | 189 | 0.58 |
|  |  |  | 9 |  |  | 98 | 1 |
|  |  |  | 10 |  |  | 275 | 0.5 |
|  |  |  | 11 |  |  | 91 | 0.71 |
|  |  |  | 12 |  |  | 131 | 0.77 |
|  |  |  | 13 |  |  | **104** | **0.6** |
|  |  |  | 14 |  |  | **197** | **0.26** |
|  |  |  | 15 |  |  | **186** | **0.21** |
|  |  |  | 16 |  |  | **216** | **0.4** |
|  |  |  | 17 |  |  | **169** | **0.6** |
|  |  |  | 18 |  |  | **213** | **0.14** |
|  |  |  | 19 |  |  | **194** | **0.5** |
|  |  |  | 20 |  |  | **104** | **0.41** |
|  |  |  | 21 |  |  | **252** | **0.24** |
|  |  |  | 22 |  |  | **262** | **0.42** |
|  |  |  | 23 |  |  | **189** | **0.21** |
|  |  |  | 24 |  |  | **141** | **0.6** |
|  |  |  | 25 |  |  | **172** | **0.7** |
|  |  |  | 26 |  |  | **144** | **0.43** |
|  |  |  | 27 |  |  | **188** | **0.67** |
|  |  |  | 28 |  |  | **49** | **0.73** |
| **2023-2024** | **Autumn** | **102** | 1 | **Natural magnetic field** | **Full-spectrum light (indoors, UV + White LEDs)** | **341** | **0.6** |
|  |  |  | 2 |  |  | **161** | **0.5** |
|  |  |  | 3 |  |  | **30** | **0.47** |
|  |  |  | 4 |  |  | **9** | **0.18** |
|  |  |  | 5 |  |  | **358** | **0.55** |
|  |  |  | 6 |  |  | **335** | **0.53** |
|  |  |  | 7 |  |  | **337** | **0.55** |
|  |  |  | 8 |  |  | **22** | **0.4** |
|  |  |  | 9 |  |  | **199** | **0.18** |
|  |  |  | 10 |  |  | **211** | **0.17** |
|  |  |  | 11 |  |  | **236** | **0.2** |
|  |  |  | 12 |  |  | **209** | **0.34** |
|  |  |  | 13 |  |  | **255** | **0.15** |
|  |  |  | 14 |  |  | **231** | **0.21** |
|  |  |  | 15 |  |  | **130** | **0.37** |
|  |  |  | 16 |  |  | **156** | **0.7** |
|  |  |  | 17 |  |  | **310** | **0.84** |
|  |  |  | 18 |  |  | **68** | **0.34** |
|  |  |  | 19 |  |  | 36 | 0.99 |
|  |  |  | 20 |  |  | 355 | 0.84 |
|  |  |  | 21 |  |  | 355 | 0.29 |
|  |  |  | 22 |  |  | 47 | 0.25 |
|  |  |  | 23 |  |  | 309 | 0.32 |
|  |  |  | 24 |  |  | 190 | 0.4 |
|  |  |  | 25 |  |  | 67 | 0.44 |
|  |  |  | 26 |  |  | 40 | 0.21 |
|  |  |  | 27 |  |  | 82 | 0.42 |
|  |  |  | 28 |  |  | 25 | 0.89 |
|  |  |  | 29 |  |  | 304 | 0.22 |
|  |  |  | 30 |  |  | 96 | 0.47 |
|  |  |  | 31 |  |  | 181 | 0.3 |
|  |  |  | 32 |  |  | 69 | 0.56 |
|  |  |  | 33 |  |  | 325 | 0.43 |
|  |  |  | 34 |  |  | 197 | 0.48 |
|  |  |  | 35 |  |  | 352 | 0.67 |
| **2024** | **Autumn** | **35** | 1 | **Natural magnetic field** | **Full-spectrum light (outdoors, no sun - diffusor)** | 42 | 0.67 |
|  |  |  | 2 |  |  | 30 | 0.74 |
|  |  |  | 3 |  |  | 196 | 0.19 |
|  |  |  | 4 |  |  | 159 | 0.52 |
|  |  |  | 5 |  |  | 155 | 0.55 |
|  |  |  | 6 |  |  | 275 | 0.4 |
|  |  |  | 7 |  |  | 244 | 0.69 |
|  |  |  | 8 |  |  | 125 | 0.78 |
|  |  |  | 9 |  |  | 52 | 0.31 |
|  |  |  | 10 |  |  | 38 | 0.98 |
|  |  |  | 11 |  |  | 94 | 0.86 |
|  |  |  | 12 |  |  | 303 | 0.7 |
| **2023** | **Autumn** | **51** | **1** | **Artificial magnetic field (-70 deg inclination)** | **Full-spectrum light (indoors, UV + White LEDs)** | 255 | 0.24 |
|  |  |  | 2 |  |  | 288 | 0.57 |
|  |  |  | 3 |  |  | 7 | 0.6 |
|  |  |  | 4 |  |  | 45 | 0.25 |
|  |  |  | 5 |  |  | 336 | 0.86 |
|  |  |  | 6 |  |  | 237 | 0.92 |
|  |  |  | 7 |  |  | 110 | 0.2 |
|  |  |  | 8 |  |  | 285 | 0.21 |
|  |  |  | 9 |  |  | 75 | 0.79 |
|  |  |  | 10 |  |  | 101 | 0.92 |
|  |  |  | 11 |  |  | 44 | 0.28 |
|  |  |  | 12 |  |  | 22 | 0.2 |
|  |  |  | 13 |  |  | 332 | 0.91 |
|  |  |  | 14 |  |  | 181 | 0.35 |
|  |  |  | 15 |  |  | 145 | 0.94 |
|  |  |  | 16 |  |  | 238 | 0.37 |
|  |  |  | 17 |  |  | 172 | 0.61 |

**Figure S3. Results of the comparison of the proportion of good vs. bad flights between different experimental groups (Fisher's exact test).**

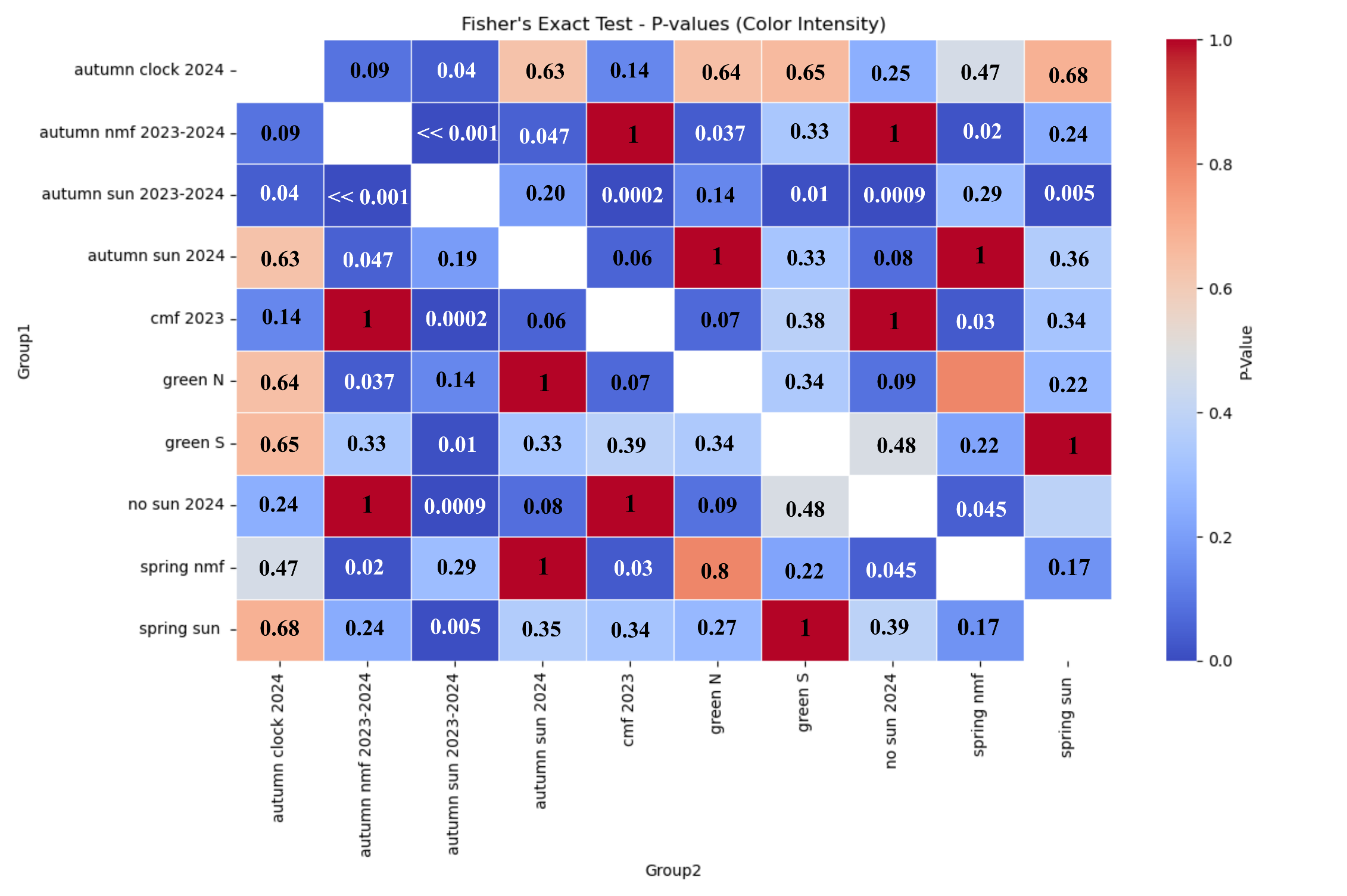

**Table S2. The results of maximum likelihood estimation for all experimental groups.**

The best fitted models are listed in corresponding column (model names are from Fitak & Johnsen, 2017)

*** -** Best model to describe distribution, **bold –** other probable models to describe distribution (deltaAICc < 2)

**A) Sun/NMF, outdoor, spring 2024**

| Model | Description | deltaAICc | AIC_weights |
| --- | --- | --- | --- |
| **M4B** | **Axial bimodal** | **0.000*** | **0.31*** |
| M5B | Bimodal | 2.38 | 0.17 |
| **M4A** | **Homogenous axial bimodal** | ***0.743*** | ***0.14*** |
| **M3A** | **Homogenous symmetric bimodal** | ***0.156*** | ***0.14*** |
| **M3B** | **Symmetric bimodal** | ***0.757*** | ***0.07*** |
| M5A | Homogenous bimodal | 3.224 | 0.06 |
| M2B | Symmetric modified unimodal | 4.110 | 0.02 |
| M2C | Modified unimodal | 5.432 | 0.013 |
| M1 | Uniform | 5.023 | 0.009 |
| M2A | Unimodal | 7.312 | 0.003 |

**B) Sun/NMF, outdoor, autumn 2023-2024**

| Model | Description | deltaAICc | AIC_weights |
| --- | --- | --- | --- |
| **M2A** | **Unimodal** | **0.000*** | **0.61*** |
| M5A | Homogenous bimodal | 3.523 | 0.19 |
| M5B | Bimodal | 6.426 | 0.08 |
| M2C | Modified unimodal | 4.833 | 0.07 |
| M2B | Symmetric modified unimodal | 5.617 | 0.04 |
| M3B | Symmetric bimodal | 8.292 | 0.012 |
| M4B | Axial bimodal | 13.044 | 0.0017 |
| M4A | Homogenous axial bimodal | 13.658 | 0.0009 |
| M1 | Uniform | 16.804 | 0.00011 |
| M3A | Homogenous symmetric bimodal | 21.075 | 0.00002 |

**C) No_Sun(diffusor)/NMF, outdoor, autumn 2024**

| Model | Description | deltaAICc | AIC_weights |
| --- | --- | --- | --- |
| **M1** | **Uniform** | **0.000*** | **0.35*** |
| M3B | Symmetric bimodal | 4.163 | 0.2 |
| M4B | Axial bimodal | 7.084 | 0.18 |
| M2B | Symmetric modified unimodal | 4.535 | 0.07 |
| M2A | Unimodal | 4.539 | 0.07 |
| M3A | Homogenous symmetric bimodal | 4.931 | 0.057 |
| M2C | Modified unimodal | 7.578 | 0.035 |
| M4A | Homogenous axial bimodal | 8.040 | 0.028 |
| M5A | Homogenous bimodal | 12.624 | 0.01 |
| M5B | Bimodal | 19.348 | 0.003 |

**D) UV-White LED/NMF, indoor, spring 2024**

| Model | Description | deltaAICc | AIC_weights |
| --- | --- | --- | --- |
| **M3A** | **Homogenous symmetric bimodal** | **0.014*** | **0.31*** |
| **M1** | **Uniform** | **0.000*** | **0.2*** |
| M4A | Homogenous axial bimodal | 2.669 | 0.14 |
| M3B | Symmetric bimodal | 2.988 | 0.11 |
| M4B | Axial bimodal | 5.597 | 0.066 |
| M5A | Homogenous bimodal | 5.942 | 0.055 |
| M2B | Symmetric modified unimodal | 4.298 | 0.036 |
| M5B | Bimodal | 9.427 | 0.03 |
| M2A | Unimodal | 4.857 | 0.03 |
| M2C | Modified unimodal | 6.242 | 0.02 |

**E) UV-White LED/NMF, indoor, autumn 2023-2024**

| Model | Description | deltaAICc | AIC_weights |
| --- | --- | --- | --- |
| **M4A** | **Homogenous axial bimodal** | **0.000*** | **0.25*** |
| **M3A** | **Homogenous symmetric bimodal** | **0.359** | **0.17** |
| **M4B** | **Axial bimodal** | **1.673** | **0.15** |
| **M3B** | **Symmetric bimodal** | **1.822** | **0.01** |
| M5A | Homogenous bimodal | 2.500 | 0.096 |
| M5B | Bimodal | 4.304 | 0.056 |
| M2B | Symmetric modified unimodal | 2.668 | 0.055 |
| M1 | Uniform | 2.418 | 0.052 |
| M2C | Modified unimodal | 3.888 | 0.036 |
| M2A | Unimodal | 3.970 | 0.028 |

**F) UV-White LED/-70 deg CMF, indoor, autumn 2023**

| Model | Description | deltaAICc | AIC_weights |
| --- | --- | --- | --- |
| **M1** | **Uniform** | **0.000*** | **0.61*** |
| M2A | Unimodal | 4.660 | 0.09 |
| M2B | Symmetric modified unimodal | 4.667 | 0.09 |
| M3A | Homogenous symmetric bimodal | 4.713 | 0.088 |
| M3B | Symmetric bimodal | 7.644 | 0.033 |
| M4A | Homogenous axial bimodal | 7.644 | 0.033 |
| M2C | Modified unimodal | 7.888 | 0.029 |
| M5A | Homogenous bimodal | 10.979 | 0.013 |
| M4B | Axial bimodal | 11.698 | 0.009 |
| M5B | Bimodal | 15.608 | 0.004 |

**G) GreenLed_North/NMF, indoor, autumn 2024**

| Model | Description | deltaAICc | AIC_weights |
| --- | --- | --- | --- |
| **M2A** | **Unimodal** | **0.000*** | **0.43*** |
| M2B | Symmetric modified unimodal | 2.292 | 0.14 |
| M5A | Homogenous bimodal | 2.969 | 0.098 |
| M5B | Bimodal | 3.025 | 0.0955 |
| M2C | Modified unimodal | 3.029 | 0.00953 |
| M1 | Uniform | 4.061 | 0.056 |
| M3B | Symmetric bimodal | 5.636 | 0.026 |
| M4A | Homogenous axial bimodal | 5.734 | 0.025 |
| M4B | Axial bimodal | 6.124 | 0.02 |
| M3A | Homogenous symmetric bimodal | 7.170 | 0.01 |

**H) GreenLed_SouthEast/NMF, indoor, autumn 2024**

| Model | Description | deltaAICc | AIC_weights |
| --- | --- | --- | --- |
| **M5A** | **Homogenous bimodal** | **0.000*** | **0.57*** |
| M5B | Bimodal | 4.074 | 0.21 |
| **M2A** | **Unimodal** | **1.164** | **0.09** |
| M2B | Symmetric modified unimodal | 2.172 | 0.06 |
| M2C | Modified unimodal | 5.081 | 0.021 |
| M3B | Symmetric bimodal | 5.241 | 0.02 |
| M4A | Homogenous axial bimodal | 6.719 | 0.009 |
| M4B | Axial bimodal | 8.631 | 0.007 |
| M1 | Uniform | 5.643 | 0.006 |
| M3A | Homogenous symmetric bimodal | 6.853 | 0.005 |

**J) Sun/NMF, outdoor, autumn 2024**

| Model | Description | deltaAICc | AIC_weights |
| --- | --- | --- | --- |
| **M2A** | **Unimodal** | **0.000*** | **0.48*** |
| M2B | Symmetric modified unimodal | 2.259 | 0.16 |
| M5A | Homogenous bimodal | 5.518 | 0.12 |
| M2C | Modified unimodal | 3.901 | 0.12 |
| M3B | Symmetric bimodal | 5.417 | 0.06 |
| M5B | Bimodal | 9.673 | 0.05 |
| M4A | Homogenous axial bimodal | 7.758 | 0.02 |
| M4B | Axial bimodal | 13.556 | 0.0021 |
| M1 | Uniform | 10.458 | 0.002 |
| M3A | Homogenous symmetric bimodal | 13.042 | 0.0007 |

**K) Sun/NMF, Clock shift +6h, outdoor, autumn 2024**

| Model | Description | deltaAICc | AIC_weights |
| --- | --- | --- | --- |
| **M2A** | **Unimodal** | **0.000*** | **0.31*** |
| M5B | Bimodal | 3.733 | 0.26 |
| **M2B** | **Symmetric modified unimodal** | **1.402** | **0.16** |
| M2C | Modified unimodal | 3.196 | 0.09 |
| M5A | Homogenous bimodal | 4.791 | 0.07 |
| M3B | Symmetric bimodal | 5.062 | 0.04 |
| M1 | Uniform | 4.206 | 0.027 |
| M4B | Axial bimodal | 7.060 | 0.023 |
| M4A | Homogenous axial bimodal | 6.055 | 0.022 |
| M3A | Homogenous symmetric bimodal | 8.442 | 0.005 |

**Table S4. Results of Moore’s modified Rayleigh test.**

| **Light source/ Magnetic field/season** | **Moore’s modified Rayleight test visualisation** | |
| --- | --- | --- |
| **A) Full-spectrum light (Sun)/NMF/ Spring 2024**  ****unimodal**  α = 335˚  n = 25  R* = 0.29  0.5 < p < 0.9  **===============**  **** axial**  α = 169±180°  R* = 1.3  n = 25  **0.005 < p < 0.01** | 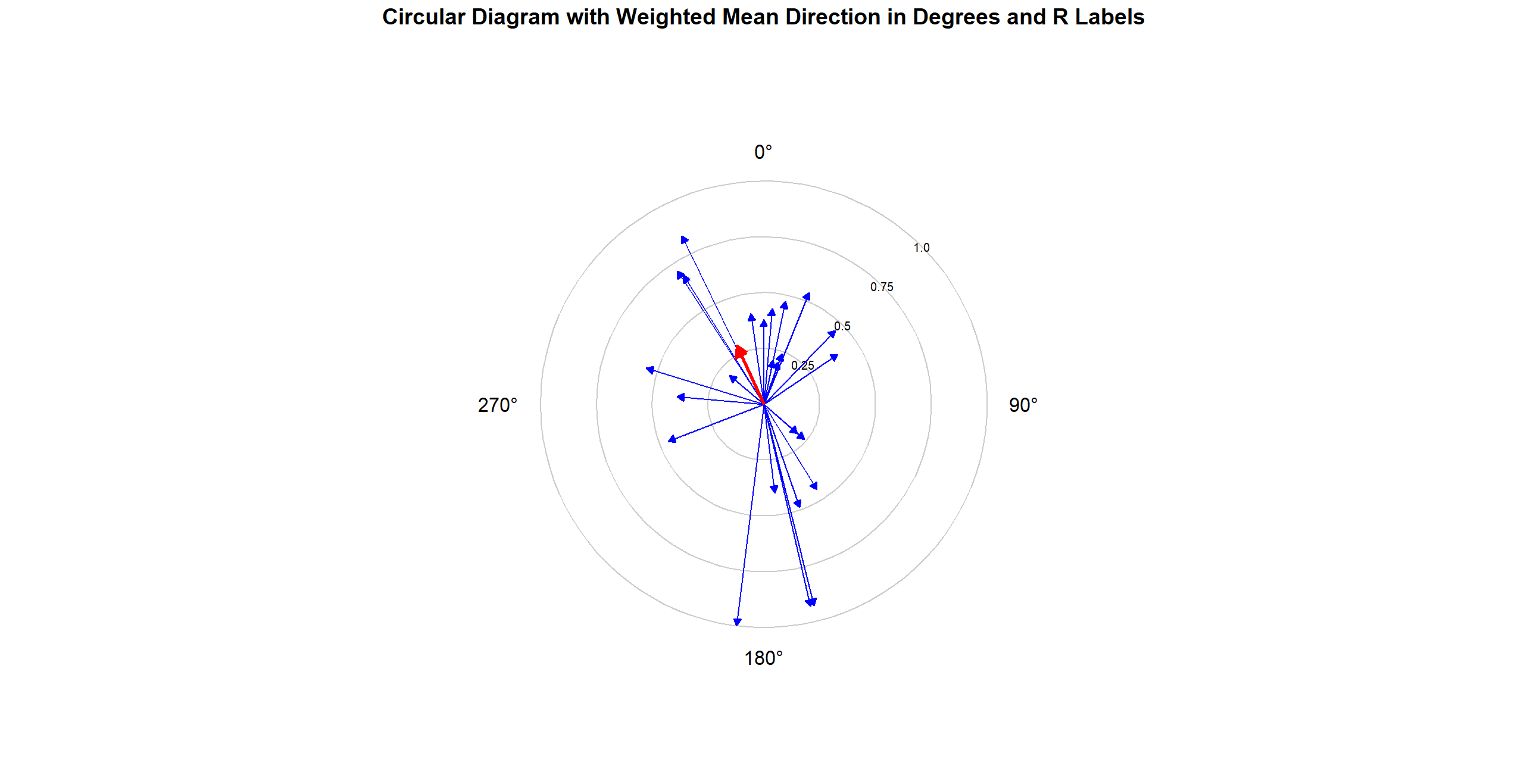 | |
| **B) Full-spectrum light (Sun)/NMF/Autumn 2023-2024**  α = 152˚  n = 28  R* = 1.62  **p < 0.001**  95% CI = 126˚ - 181˚ | 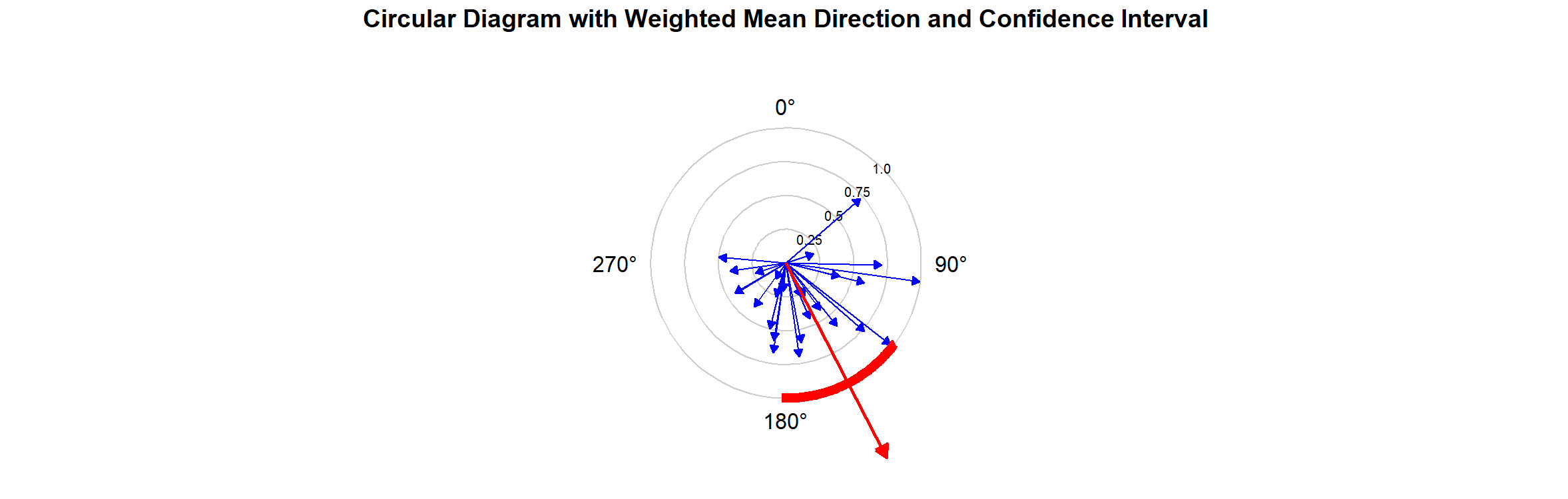 | |
| **C) No_Sun (diffisor)/NMF/ Autumn 2024**  α = 70˚  n = 12  R* = 0.61  0.1 < p < 0.5 | 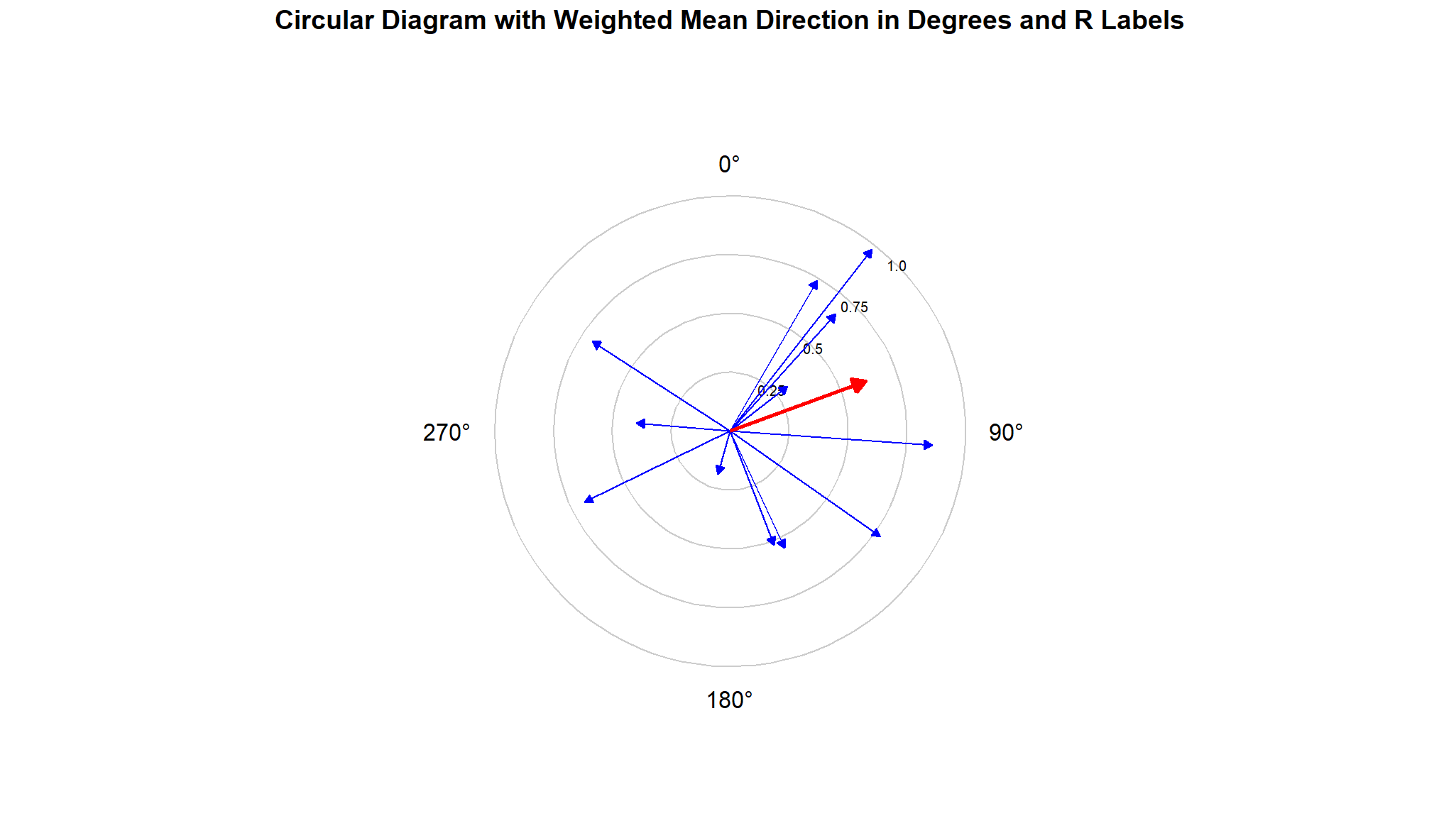 | |
| **D) UV-White LED/NMF/ Spring 2024**  **unimodal  α = 164˚  n = 17  R* = 0.58  0.1 < p < 0.5  **===============**  ****** axial  α = 179  n = 17  R* = 0.98  0.05 < p < 0.1 | 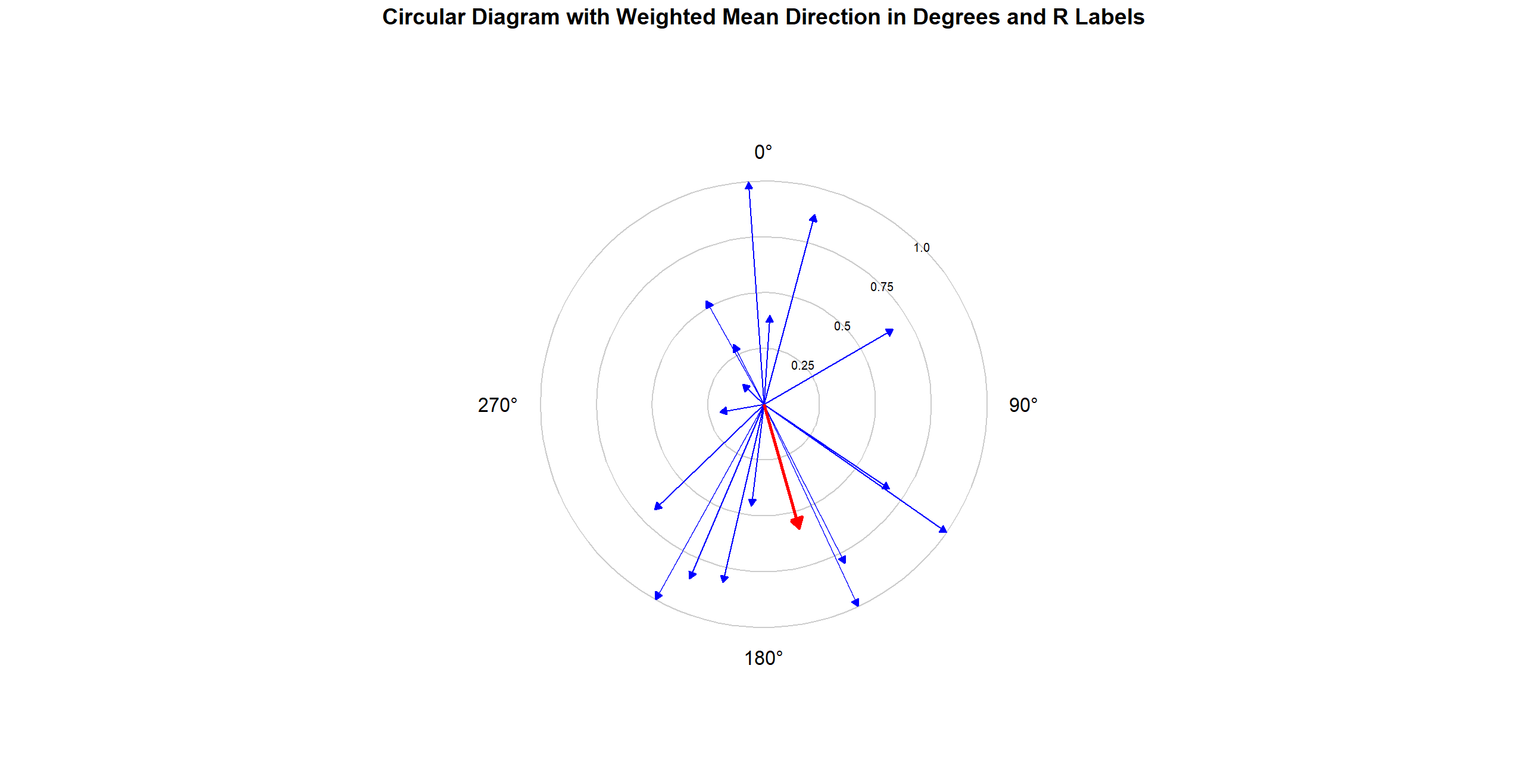 | |
| **E) UV-White LED/NMF/ Autumn 2023-2024**  α = 18˚  n = 35  R* = 1.11  **p < 0.05**  **____________**  ****** axial  α = 180  n = 35  R* = 1.04  0.025 < p < 0.05 | 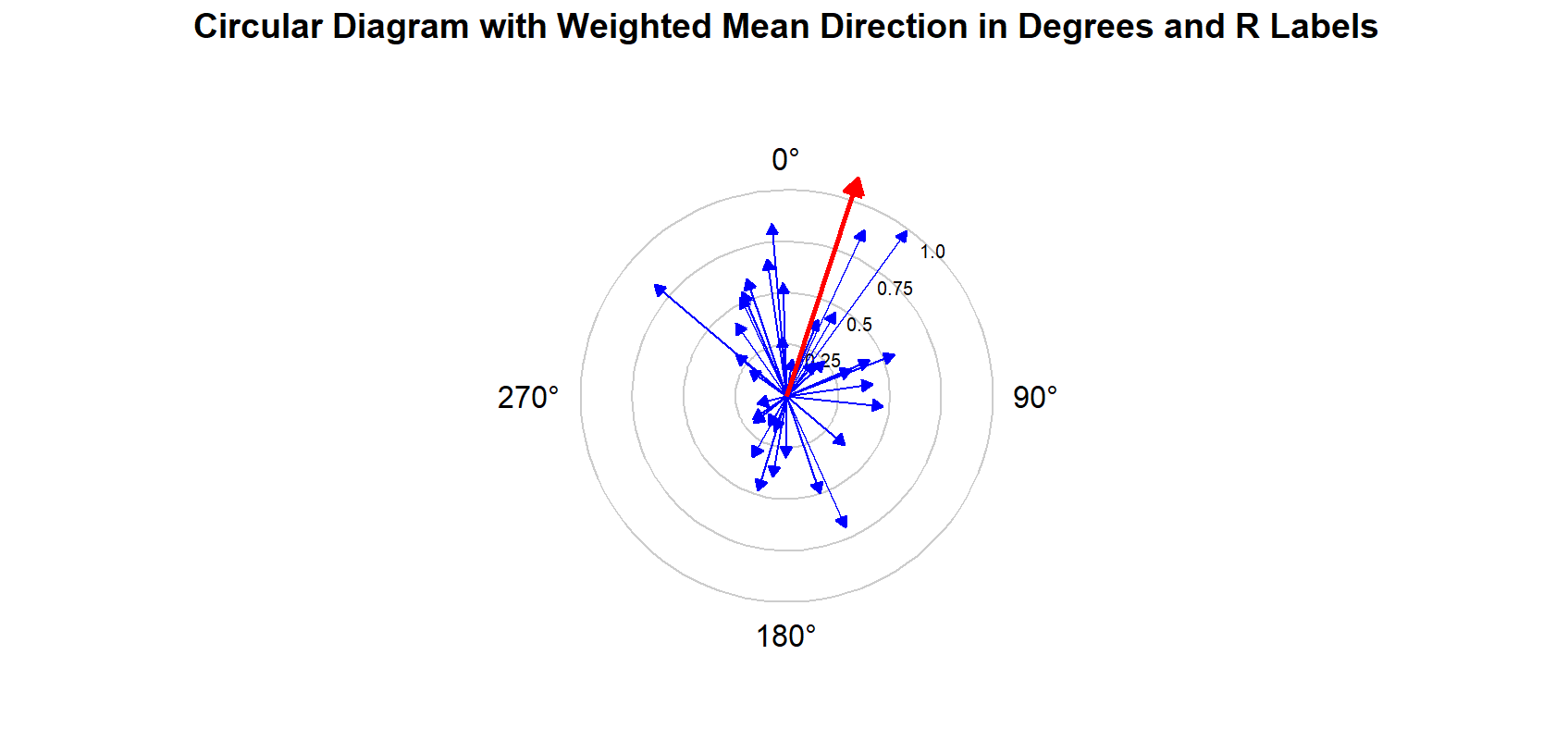 | 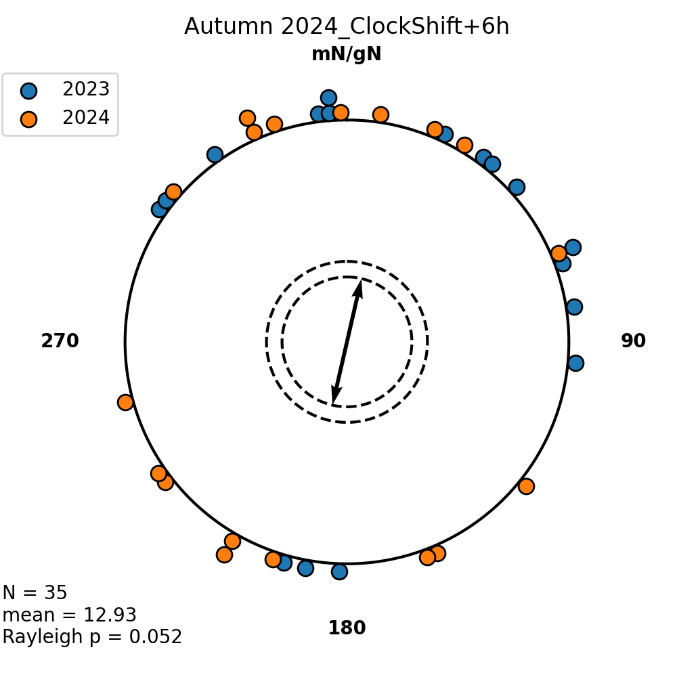 |
| **2023:**  α = 33˚  n = 17  R* = 1.04  **0.025 < p < 0.05** | 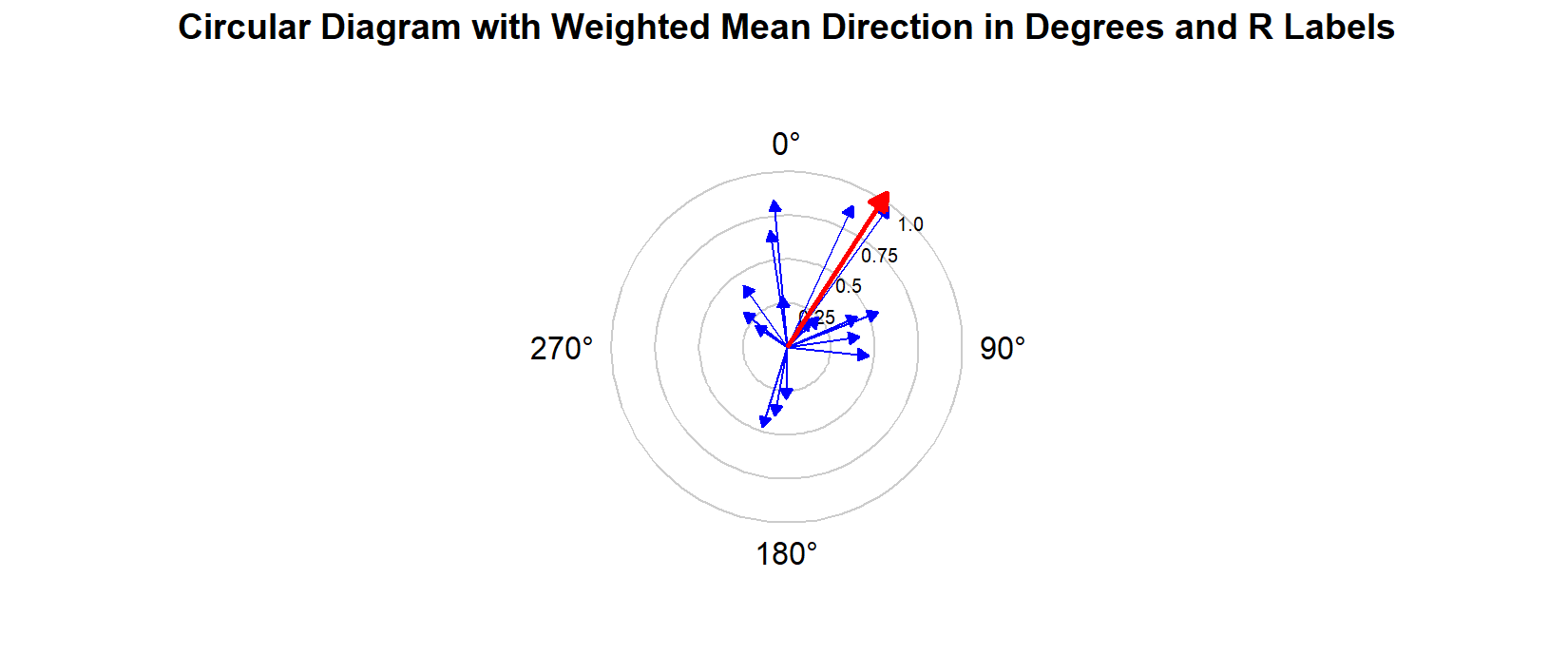 | 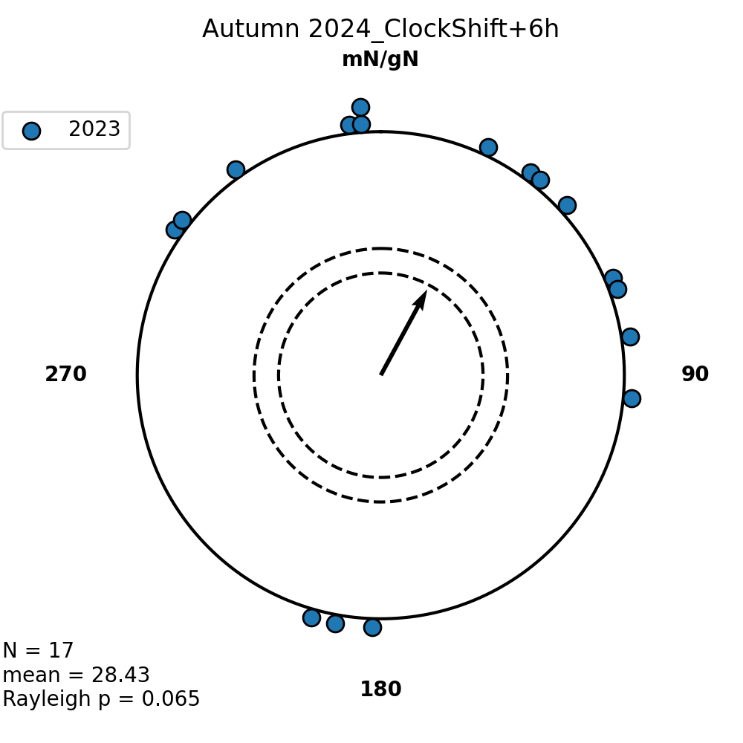 |
| **2024:**  α = 346˚  n = 18  R* = 0.57  0.1 < p < 0.5 | 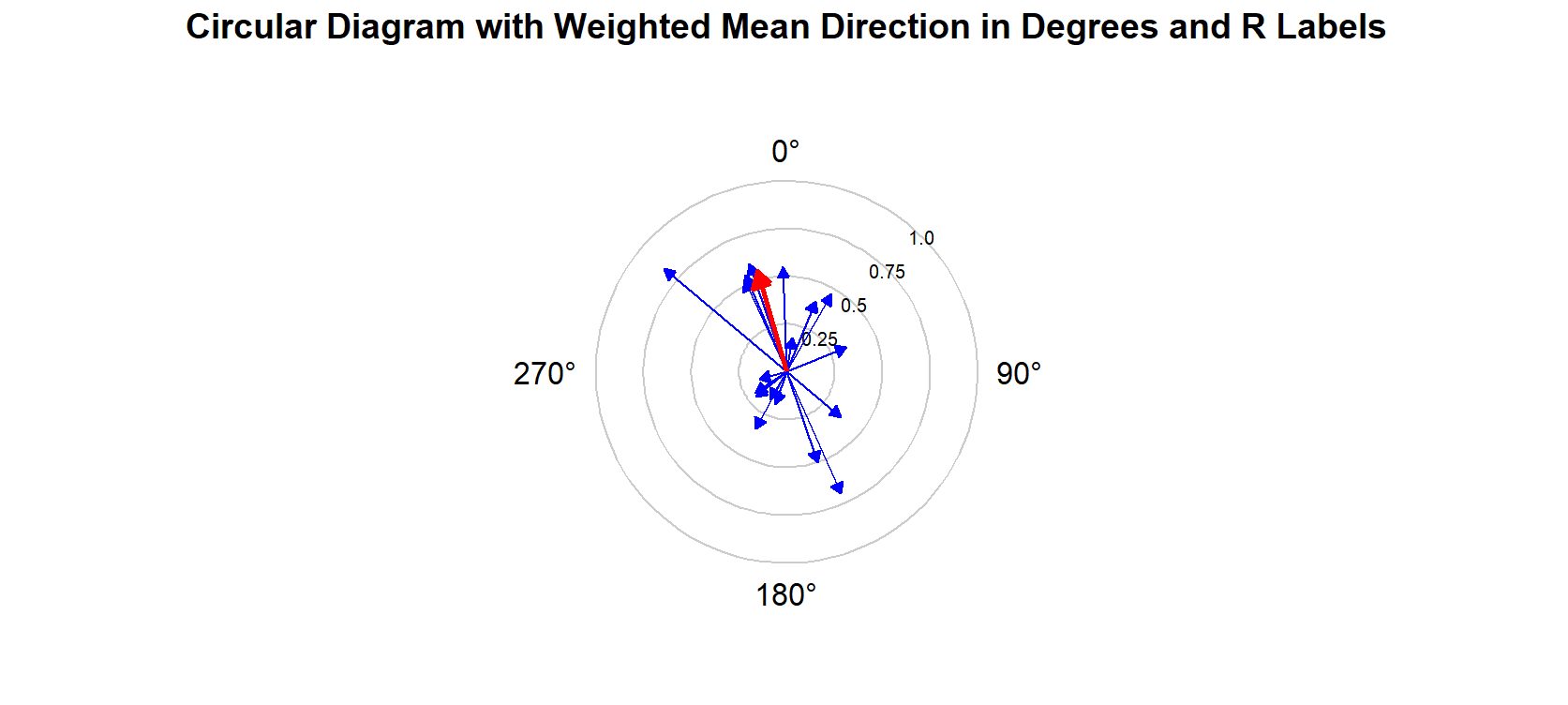 | 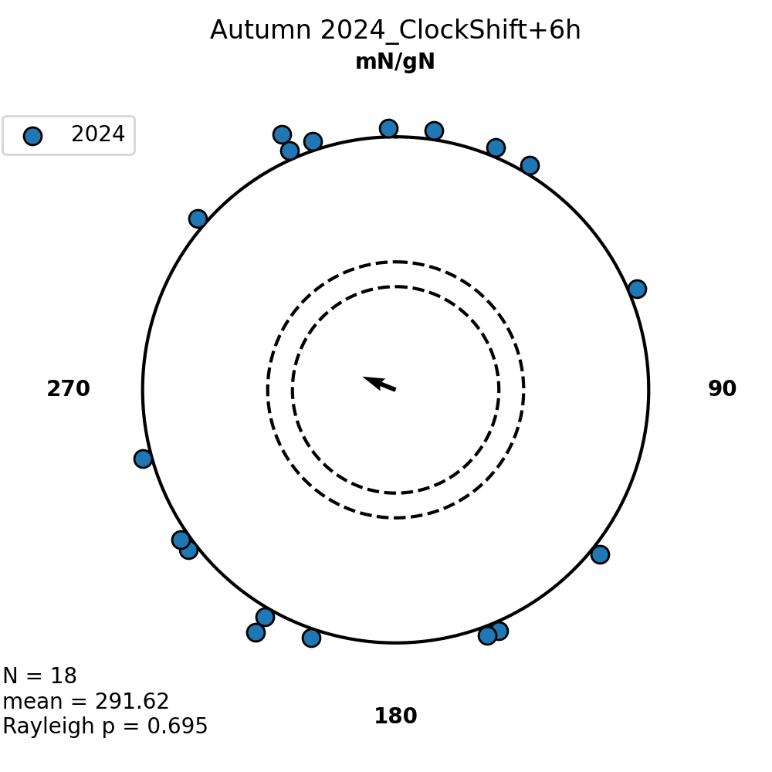 |
| **F) UV-White LED/-70 deg CMF/ Autumn 2023**  α = 53˚  n = 17  R* = 0.04  0.99 < p < 0.995 | 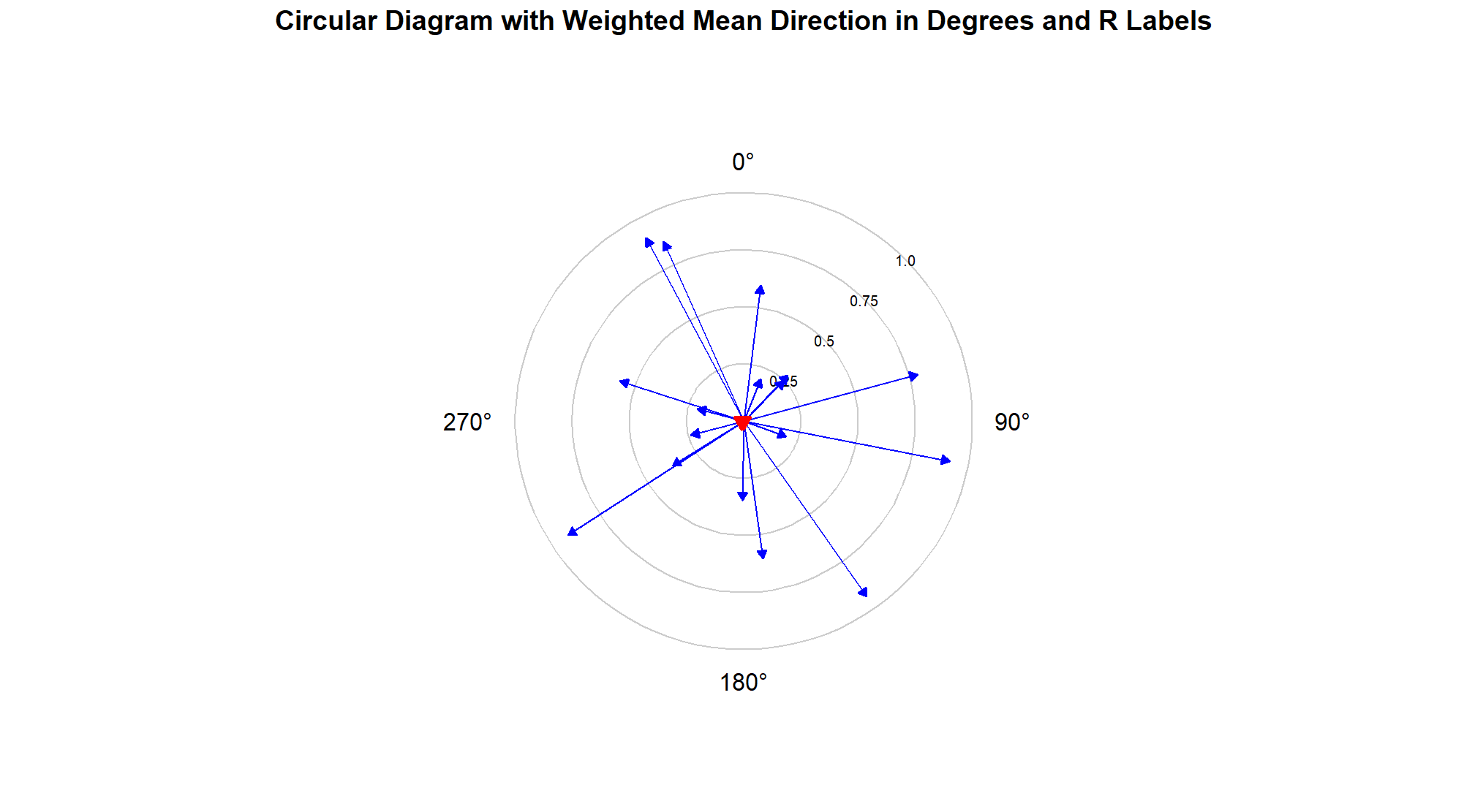 | |
| **G) GreenLed_North/NMF/ Autumn 2024**  α = 303˚  n = 18  R* = 1.022  0.05 < p < 0.1 | 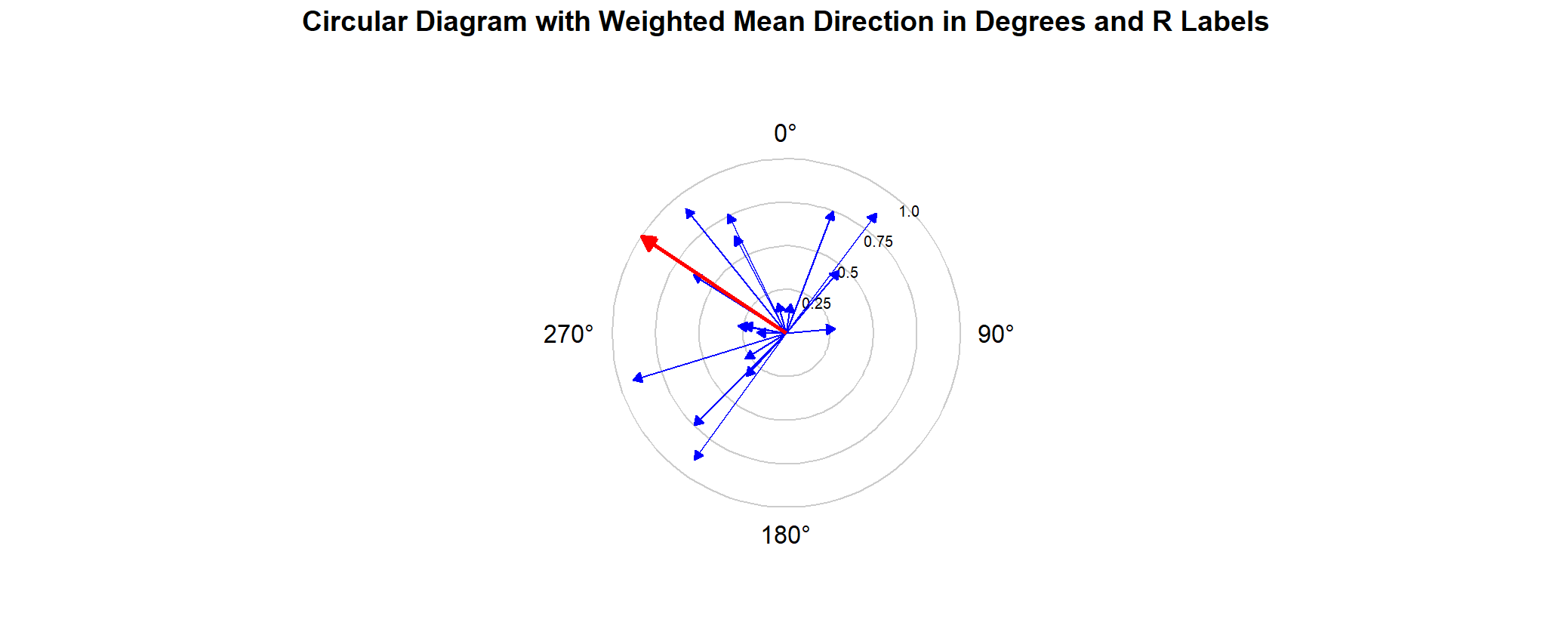 | |
| **H) GreenLed_SouthEast/**  **NMF/ Autumn 2024**  α = 45˚  n = 17  R* = 0.86  0.1 < p < 0.5 | 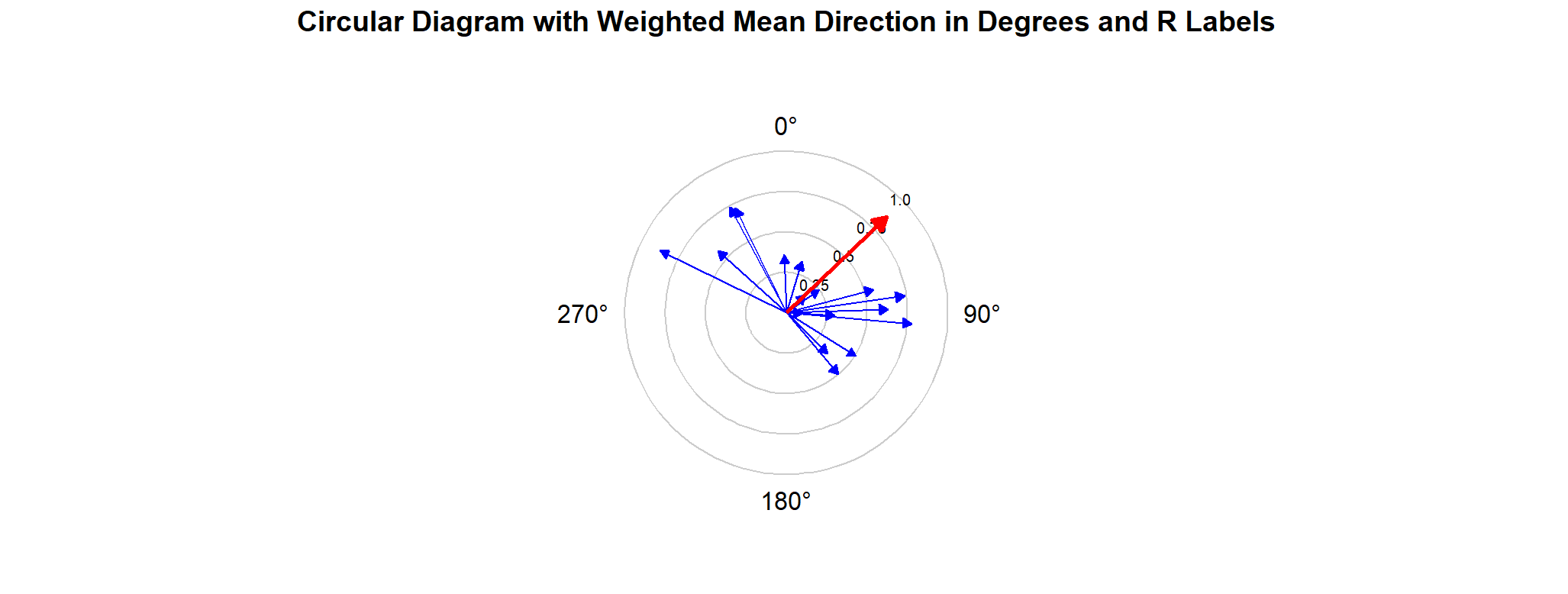 | |
| **J) Full-spectrum light (Sun)/NMF/ Autumn 2024**  α = 162˚  n = 16  R* = 1.29  **0.005 < p < 0.01**  95% CI = 121˚ - 191˚ | 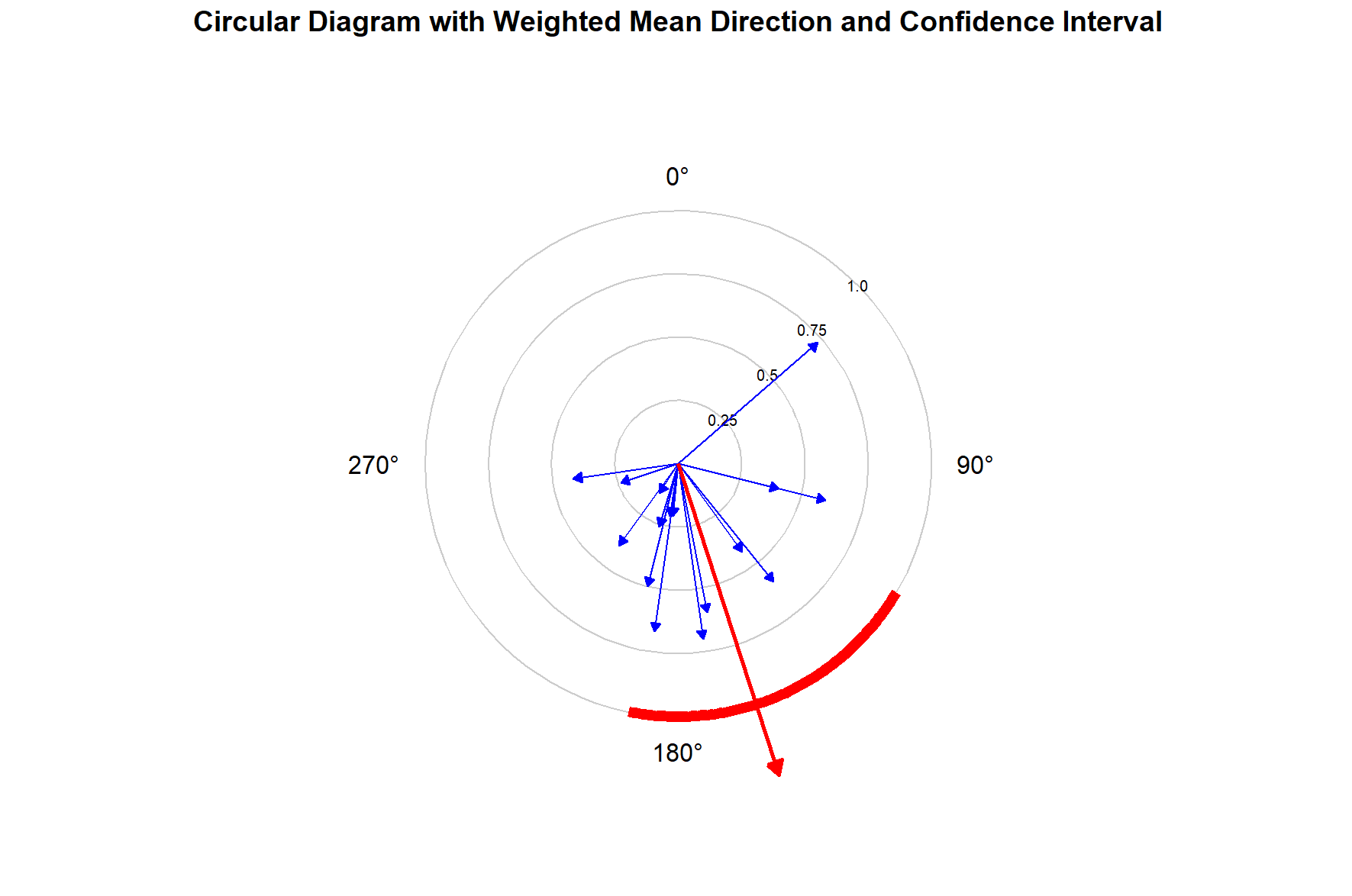 | |
| **K) Full-spectrum light (Sun)/NMF/ Clock shift +6h/Autumn 2024**  α = 195˚  n = 21  R* = 1.05  **0.025 < p < 0.05**  95% CI = 148˚ - 240˚ | 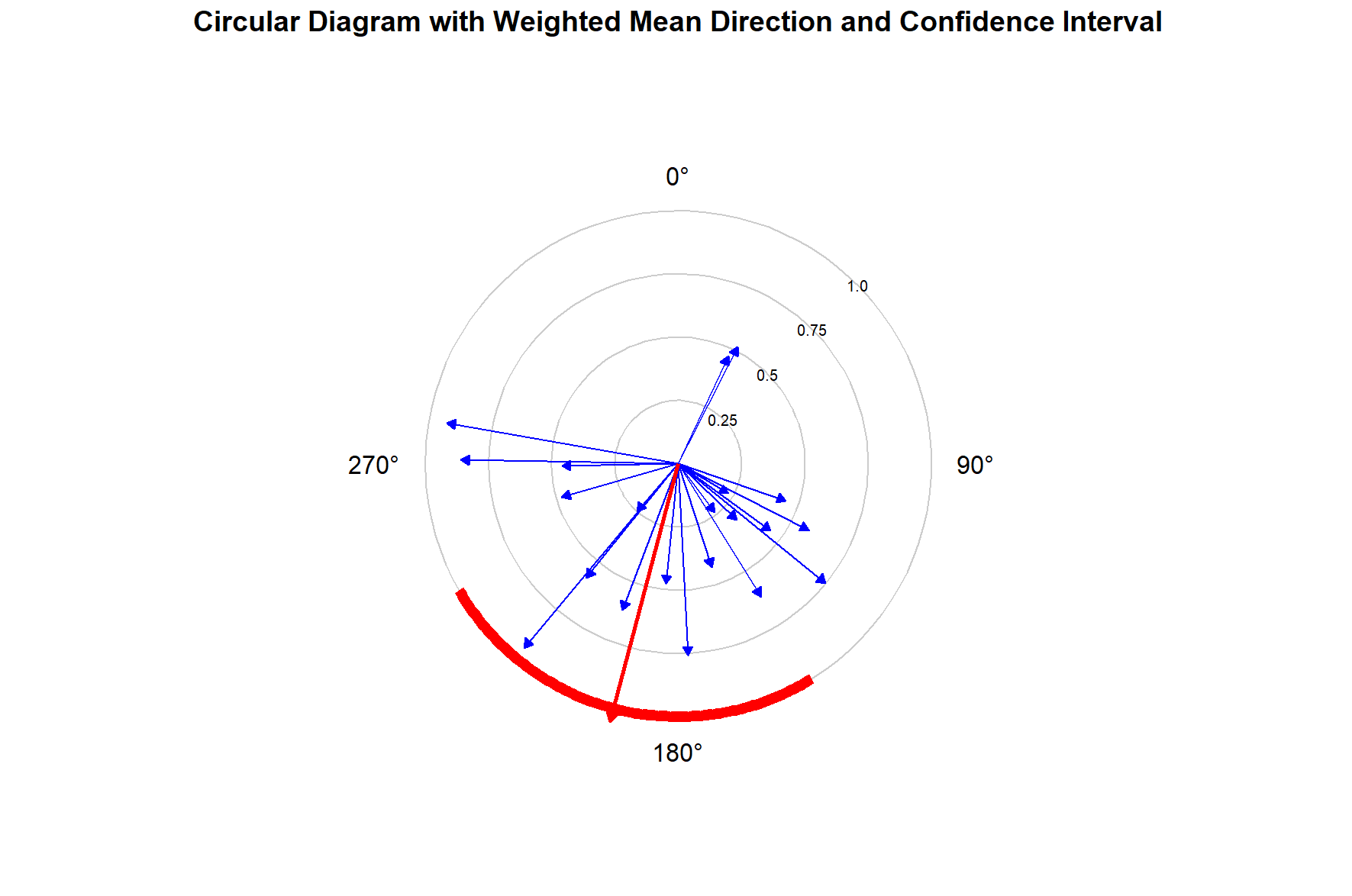 | |

**Figure S5. Vector strength (r_value between 0 and 1) in different experimental groups:

LED 1** - N_GreenLED group, **LED 2** - SE_GreenLED group**; LED 3** – UV-White NMF spring 2013 group, **LED 4** – UV-White NMF 2023-2024 autumn group; **LED 5** - -70˚CMF group; **No sun** – Overcast group; **Sun 1** – Control group autumn 2024; **Sun 2** – Clock shift group; **Sun 1** – Control group spring 2024; **Sun 4** – Control group autumn 2023-2024.

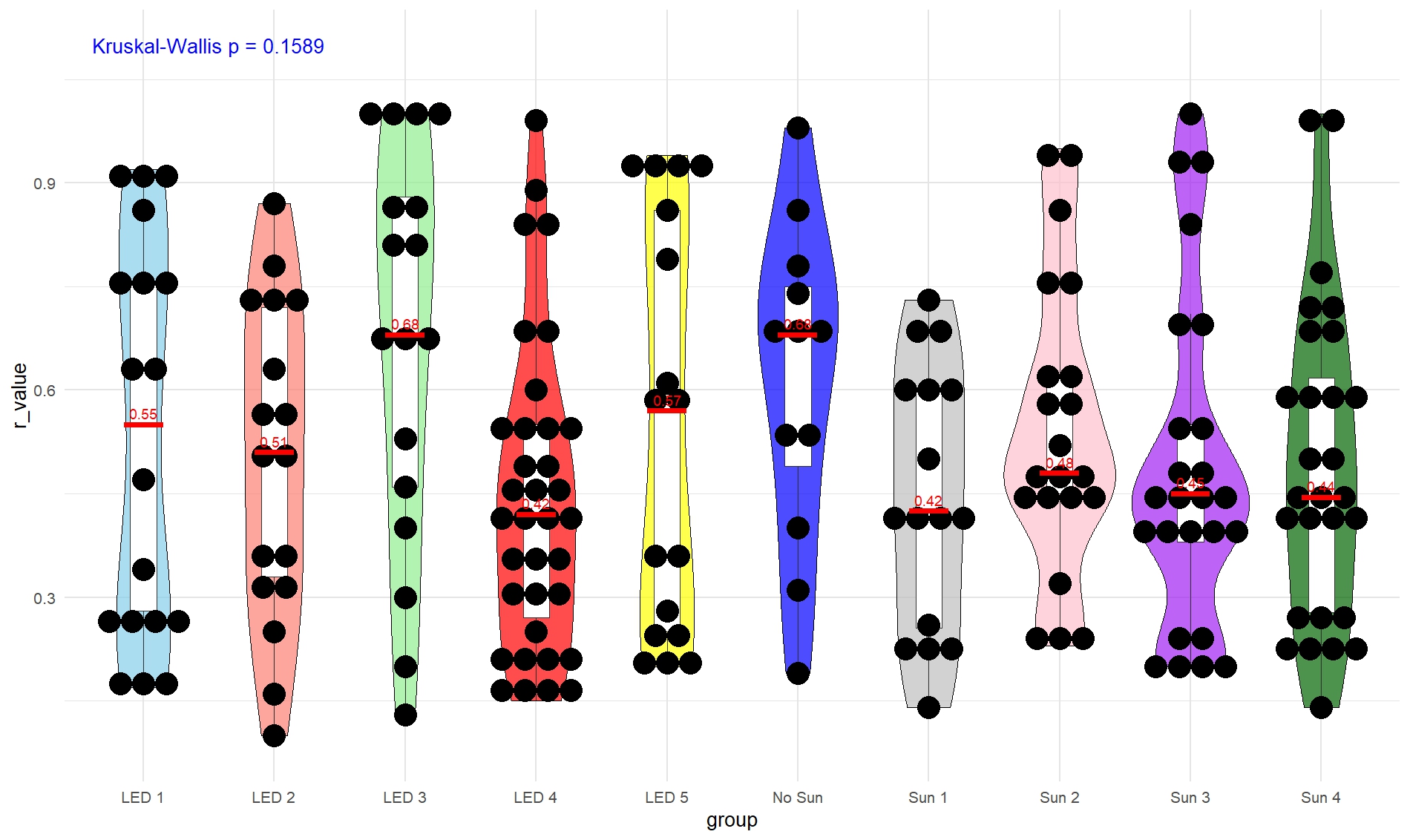

**Figure S6. Results of bootstrap analysis.**

Each diagram represents a distribution of lengths of mean vectors that were calculated using a bootstrap technique (n = 100000, see details in the main text of manuscripts, Materials and methods section): A) red admirals tested under artificial LED light conditions in the NMF indoors in 2023-2024; B) red admirals tested under overcast conditions in the NMF outdoors in 2024; C) red admirals tested under artificial LED light conditions in the -70˚CMF indoors in 2023. Vertical blue and red lines indicate 95 and 98 % quantiles, respectively. The red curve is a normal distribution, an orange dot is a length of the mean vector of group in each experimental condition

| **A** | **Sun vs Led 2023-2024**  > quantile(r, c(0.025, 0.975))  2.5% 97.5%  0.4507416 0.7329590  > quantile(r, c(0.009, 0.999))  0.9% 99.9%  0.4179058 0.8013545 | 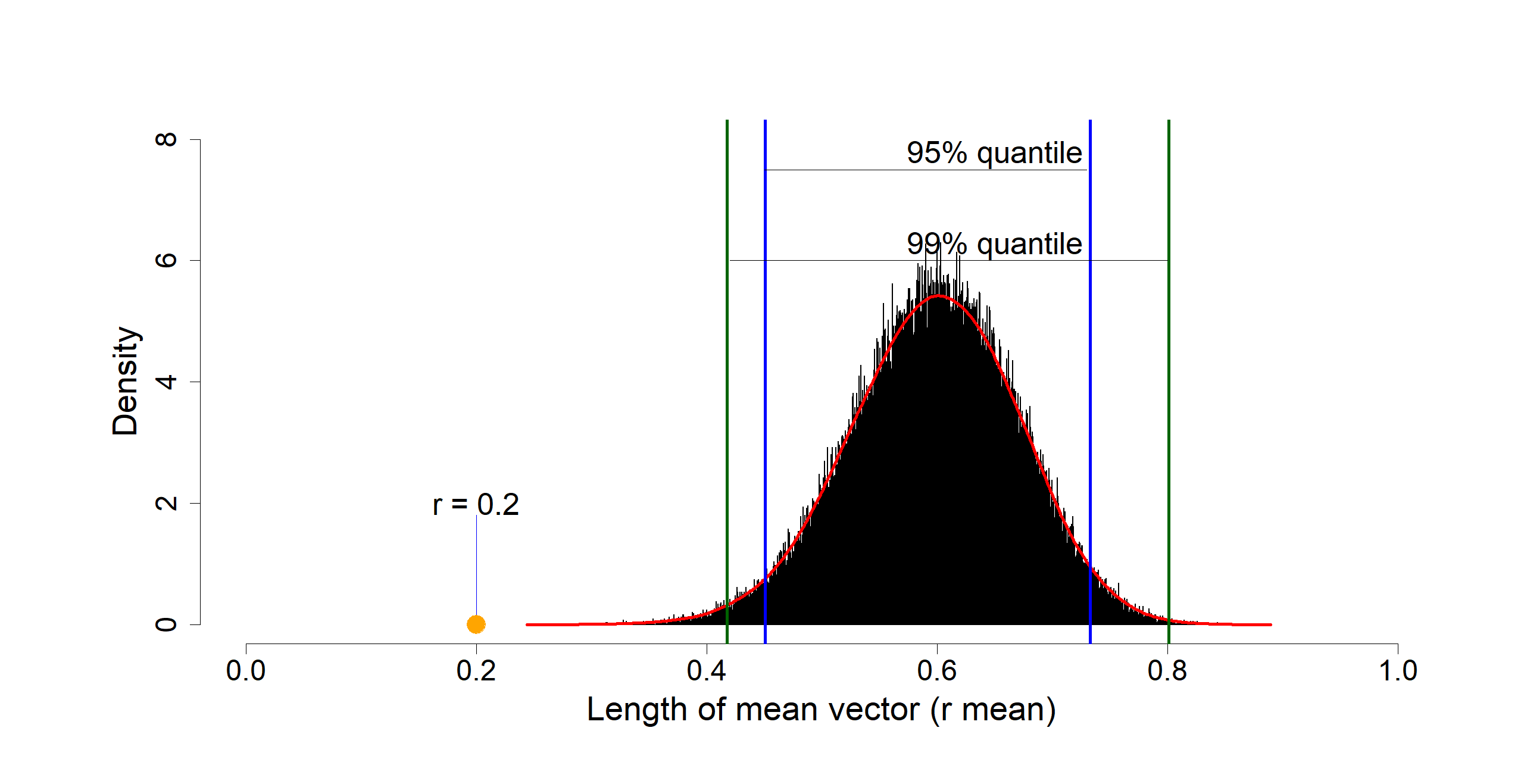 |
| --- | --- | --- |
| **B** | **Sun 2024 vs No Sun 2024**  > quantile(r, c(0.025, 0.975))  2.5% 97.5%  0.4115027 0.8890906  > quantile(r, c(0.009, 0.999))  0.9% 99.9%  0.3522340 0.9602213 | 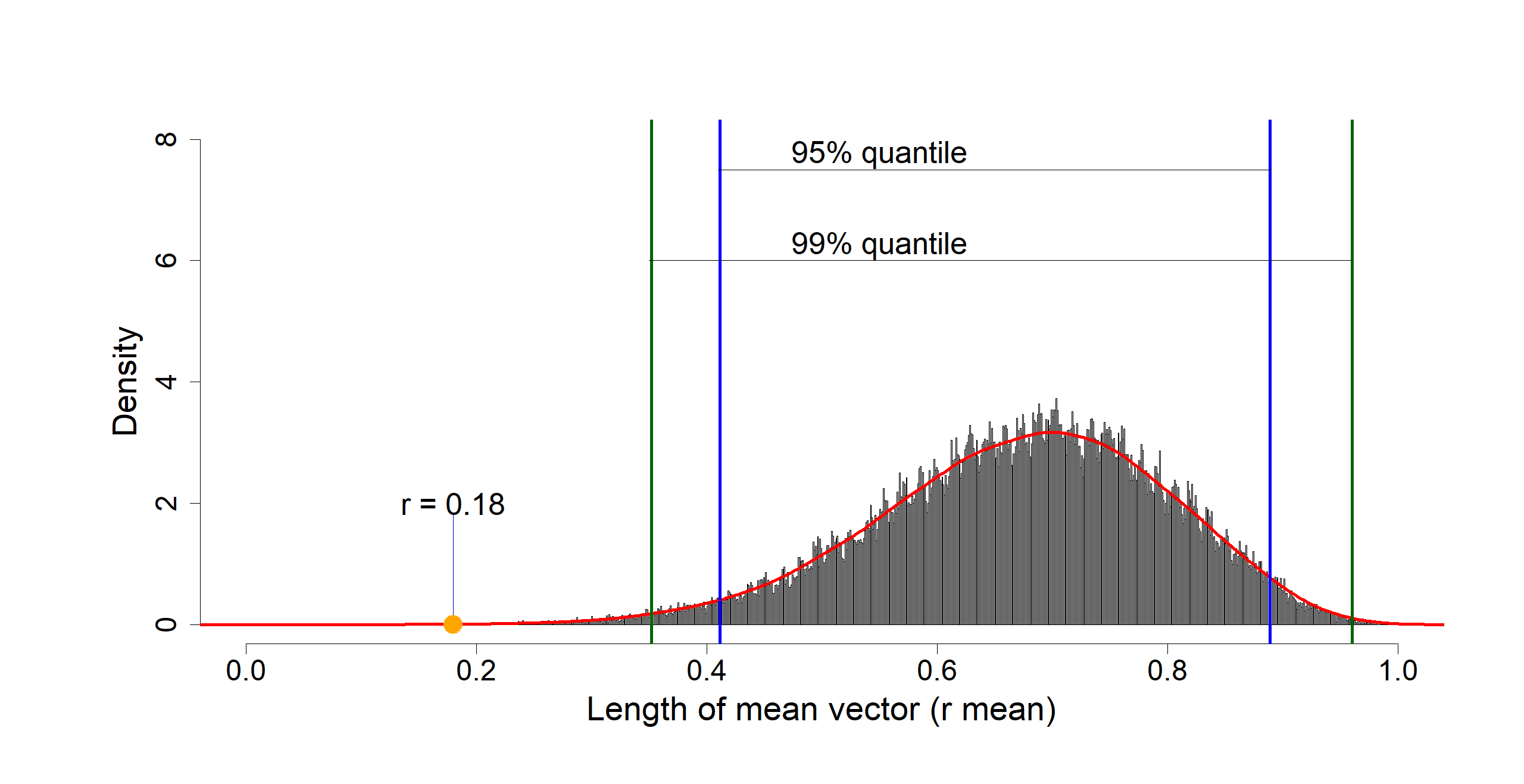 |
| **C** | **Sun 2023 vs -70 deg CMF 2023**  > quantile(r, c(0.025, 0.975))  2.5% 97.5%  0.3425620 0.7774345  > quantile(r, c(0.009, 0.999))  0.9% 99.9%  0.2962159 0.8804595 | 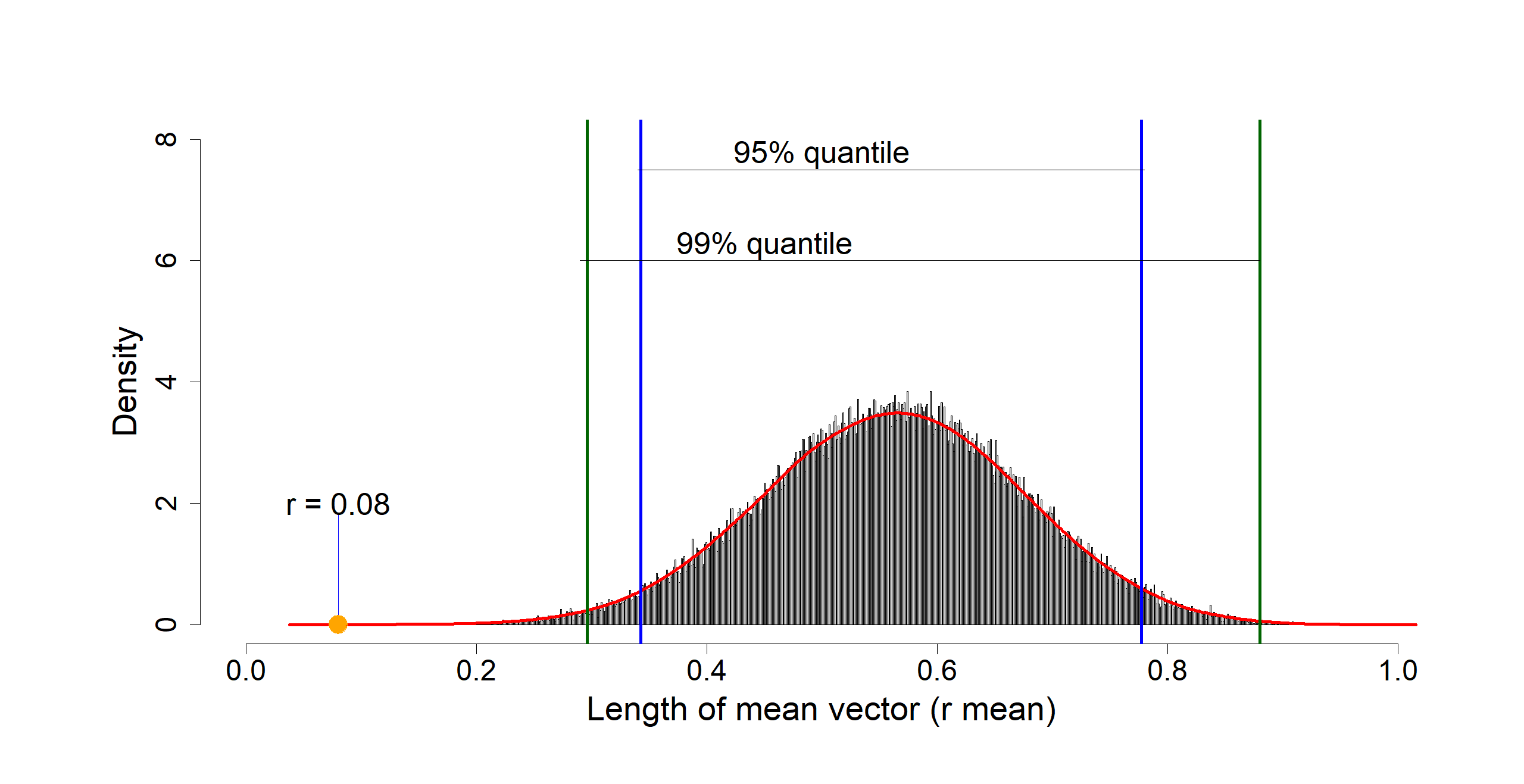 |
